# RareFoldGPCR: Agonist Design Beyond Natural Amino Acids

**DOI:** 10.1101/2025.10.01.679733

**Authors:** Qiuzhen Li, Thomas Helleday, Patrick Bryant

## Abstract

Noncanonical amino acids (NCAAs) expand the chemical diversity of peptides beyond the twenty standard residues, offering new opportunities for designing binders with novel interaction modes and functional activity. G protein-coupled receptors (GPCRs) are central to cellular signalling and represent one of the largest classes of therapeutic targets, yet their functional modulation remains challenging. Here, we present RareFoldGPCR (RFG), a GPCR-specialised AI model for structure prediction and design that supports NCAAs. By applying transfer learning on high-resolution GPCR structures, RFG accurately models and rationally designs both linear and cyclic peptides that incorporate NCAAs and modulate GPCR activity. This is achieved without the model ever being trained on NCAA-based GPCR modulators. We showcase the capability of RFG by designing peptide agonists for the glucagon-like peptide-1 receptor (GLP1R) and validating their functional activity experimentally in cell-based assays. We investigate the precise capabilities of generating active agonists by expanding different regions of the native GLP-1 hormone, and further demonstrate the design of cyclic peptide agonists with entirely novel sequences and topologies, creating new agonist modes. We analyse how design metrics relate to pathway specificity, enabling precise modulation of pathway activity, such as activating the cAMP response without recruiting β-arrestin to reduce receptor desensitisation. RFG shows how transfer learning on specific target classes enables generalisation to new chemistry and molecular topology, providing a broadly applicable strategy for designing functional ligands beyond the constraints of natural amino acid chemistry. RFG is freely available: https://github.com/patrickbryant1/RareFoldGPCR

## Introduction

G protein-coupled receptors (GPCRs) are one of the most important and widely studied classes of membrane proteins. GPCRs are central to physiological processes such as hormone sensing [1,2], neurotransmission [3,4], and immune regulation [5,6]. They are also among the most important therapeutic targets, with over one-third of FDA-approved drugs acting on GPCRs [7,8]. However, most GPCR ligands have been identified through laborious screening and optimisation; in addition, only a fraction of the receptor family has been successfully targeted [7,9]. A core limitation remains: the lack of a general, rational method to design functional molecules that modulate specific GPCRs with precision. Here, peptides are especially important as many GPCRs, e.g. the important metabolic GCGR, GLP1R and GIPR, have large orthosteric pockets unsuitable for small molecules [10]. At the same time, antibodies are too large and sterically hindered to effectively enter the orthosteric sites.

Recent advances in deep learning have enabled targeted binder design from structural templates (e.g. BindCraft [11], ProteinMPNN [12], RFDiffusion [13]). These advances build on breakthroughs in protein structure prediction, such as AlphaFold2 [14] and AlphaFold-multimer [15]. However, these methods remain intrinsically tied to structural priors and natural amino acid chemistry, limiting their ability to produce novel topologies or explore interfaces inaccessible to traditional protein scaffolds. Peptide-specific approaches have emerged to address some of these limitations. RFPeptides [16] extended diffusion-based design to cyclic peptides, while our model EvoBind2 [17] introduced a sequence-based strategy that bypasses structural input altogether. Importantly, EvoBind2’s sequence-based nature enabled a key breakthrough: the blind design of dual cyclic peptide agonists targeting GCGR and GLP1R [18]. Still, all of these models are limited to the standard 20 amino acids present in AlphaFold2 [14], restricting the chemical diversity available for peptide design.

The incorporation of noncanonical amino acids (NCAAs) expands the accessible chemical space, enabling the introduction of novel interaction motifs that may lead to the rational design of entirely new pharmaceuticals. We have now extended EvoBind2 to support NCAAs through RareFold, creating EvoBindRare, the first experimentally validated framework capable of protein structure prediction and design with 29 NCAAs [19]. These NCAAs introduce distinct chemistries, e.g. phosphate groups or large side-chains, providing access to geometries and interactions not represented in canonical amino acid proteins. This expansion to 49 building blocks represents a step-change in design capability and opens new possibilities for peptide therapeutics.

Here we introduce RareFoldGPCR (RFG), a GPCR-specialised extension of RareFold fine-tuned on all high-quality structures from GPCRdb [20]. Unlike EvoBind2 [17] or EvoBindRare [19], RFGs GPCR-specific transfer learning allows it to generalise to NCAA-GPCR agonists and produce experimentally validated designs. This shifts the scope from binder generation to functional design, opening access to previously unexplored chemical and conformational space for GPCR modulation.

## Results

### Fine-tuning RareFold for GPCR-peptide modelling

RareFold, originally trained on proteins limited to 256 residues, has limited performance on larger structures such as GPCRs. As the model cannot be directly applied to GPCR modelling, we fine-tuned RareFold using all available GPCR structures from GPCRdb [20] (Figure 1a). As RareFold handles 29 NCAAs, the ability to model 49 amino acid types is combined with the ability to model GPCRs, even though the GPCRs do not contain NCAAs. The fine-tuned single-chain model was evaluated on the orthogonal task of predicting GPCR-peptide complexes (Figures 1b and 1c). At step 500, the model achieves the lowest peptide RMSDs; however, the median value remains high (approximately 20 Å), indicating the need for further refinement. To address this, we selected 315 GPCR-peptide complexes and continued fine-tuning specifically on these interactions. Although the model remains formally single-chain, we introduced a chain break to enable learning of peptide-receptor interactions [21]. This allows for utilising single-chain models like RareFold in a transfer learning setting to learn all-atom structure prediction of GPCR-peptide complexes with high accuracy.

**Figure 1.**
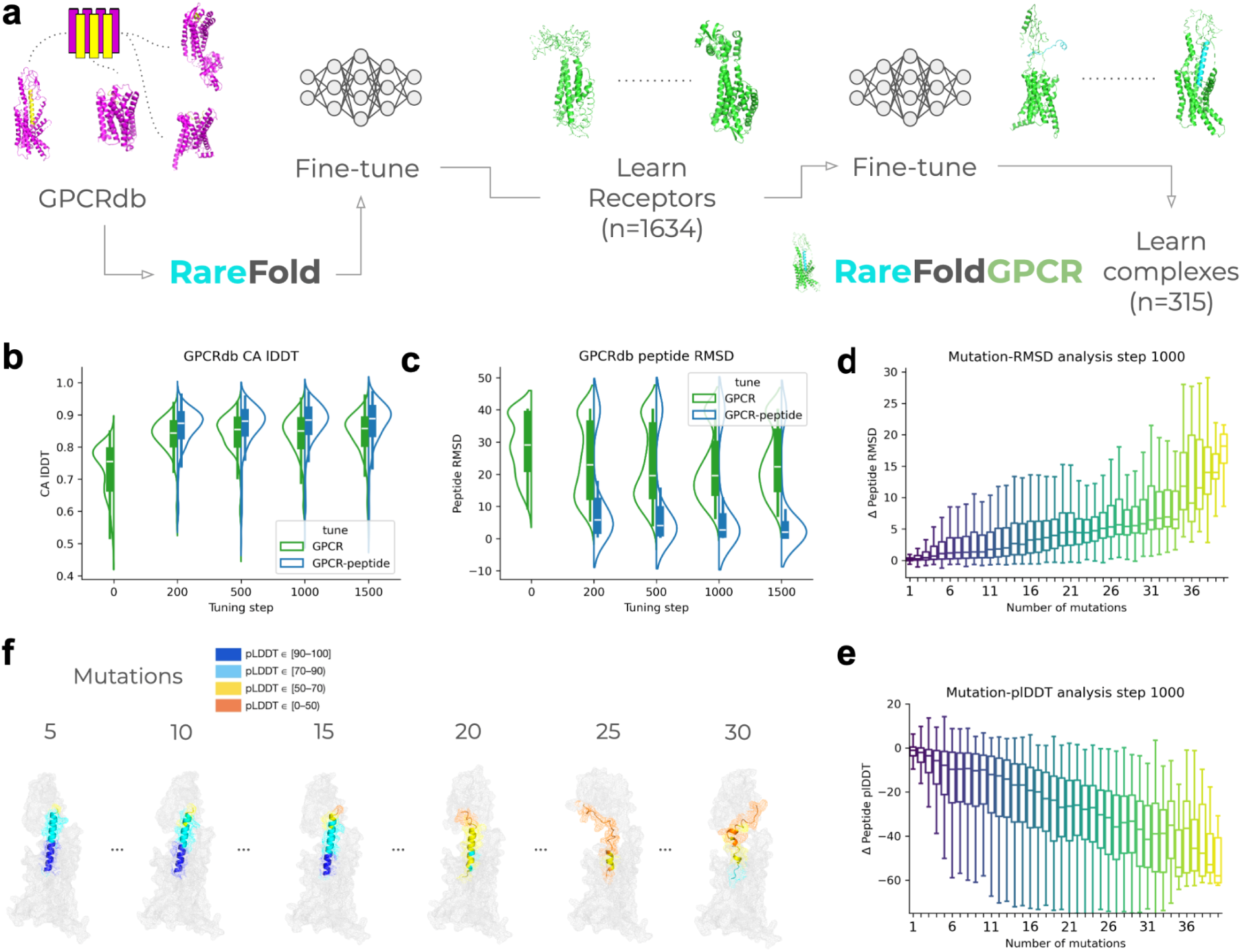
Fine-tuning RareFold for GPCR and GPCR-peptide modelling. **a)** Schematic overview of the fine-tuning process. Starting from the original RareFold model, we fine-tune on single-chain GPCR structures (GPCR sc) to improve receptor modelling, followed by further tuning on GPCR–peptide complexes to enable learning of GPCR-specific binding patterns. Each step is evaluated both on receptor structure prediction and on complex formation, including in silico mutational evaluation. **b)** Improvement in GPCR structural quality (n=314) across fine-tuning steps, measured by the Cα–lDDT score. The complex mode shows higher Cα–lDDT scores, likely due to improved residue packing and structural consistency. **c)** Peptide Cβ RMSD (capped at 40 Å) versus tuning step, highlighting improvements in peptide positioning. Analysis is limited to 242 complexes where the peptide sequences contain only the 49 amino acid types supported by RareFold. **d)** Impact of single amino acid mutations on peptide Cβ RMSD at fine-tuning step 1000 for the complex model. The plot shows the change in RMSD (Δ RMSD) as a function of the number of mutations introduced. Outliers have been excluded for clarity. Peptides that could be predicted with RMSD <2 Å (n = 126 complexes) were used for the mutational analysis (n = 24’217 mutations) and 10 mutations were sampled at random for each mutation number. **e)** Impact of mutations on peptide structural confidence at fine-tuning step 1000 for the complex model. The change in peptide plDDT (Δ plDDT) is plotted against the number of mutations. Peptides that could be predicted with RMSD <2 Å (n = 126 complexes) were used for the mutational analysis (n = 24’217 mutations), and 10 mutations were sampled at random for each mutation number. **f)** Examples of predicted peptide-receptor complexes. The receptor is shown in grey, while the designed peptides are coloured according to plDDT confidence. Each example illustrates peptides carrying different numbers of introduced mutations, highlighting how increased mutational load affects both the predicted binding mode and structural confidence.

To evaluate the effects of complex tuning beyond potential overfitting to the training complexes, we define an orthogonal assessment task focused on interface recognition rather than structure reconstruction. Inspired by our previous work with EvoBind [22,23], we analyse whether the fine-tuned model can distinguish between plausible and implausible peptide-receptor interfaces. Specifically, we introduce random mutations into the peptides of complexes for which the model achieves a peptide Cβ RMSD <2 Å, using only the 20 canonical amino acids. We then evaluate the predicted confidence (plDDT, predicted local distance difference test [14,24]) of the peptide and measure the deviation of Cβ positions from the original (native) prediction, using weights from the complex-tuned model. We focus on Cβ positions to avoid confounding effects from side chain conformations that change with mutations. This setup enables us to test whether the model has merely learned to saturate interfaces or can resolve atomic-level differences, an essential property for reliable downstream applications such as de novo binder design.

Tuning step 1000 provides a good balance between complex and receptor accuracy. At step 1500, we find that the secondary structure dissolves for some receptors, although the peptide RMSD is lower, suggesting overfitting. Therefore, we select the parameters at step 1000 for the GPCR-peptide tuning, creating RareFoldGPCR (RFG). We find that the CB RMSD of the peptide increases with the number of introduced mutations (Figure 1d), and the plDDT decreases (Figure 1e), suggesting that the network has learned to recognise specific amino acid combinations necessary for GPCR-peptide interactions (Figure 1f). Based on the model’s ability to distinguish true from mutated peptides, we invert the model and continue with designing peptides to obtain novel agonists incorporating NCAAs learned previously with RareFold.

### Peptide Binder Design with RareFoldGPCR

Since RFG can distinguish between native and mutated binders, it may also be leveraged for *de novo* binder design by inverting the network [11,19,22,25]. To test this, we selected eight GPCR-binder complexes that RFG predicts with high confidence, alongside one case where the native complex is predicted poorly (8ZH8) and another where no prediction was possible due to the presence of a non-standard amino acid not yet supported by RFG (7YON, containing TYC) (Table 1). Including these examples allowed us to assess whether high-confidence designs could be generated even when the native complex is not accurately recapitulated. This provides a direct test of whether RFG can propose alternative high-confidence solutions, suggesting novel binding modes, particularly relevant for cyclic designs, which inherently rely on non-native interactions.

**Table 1.**
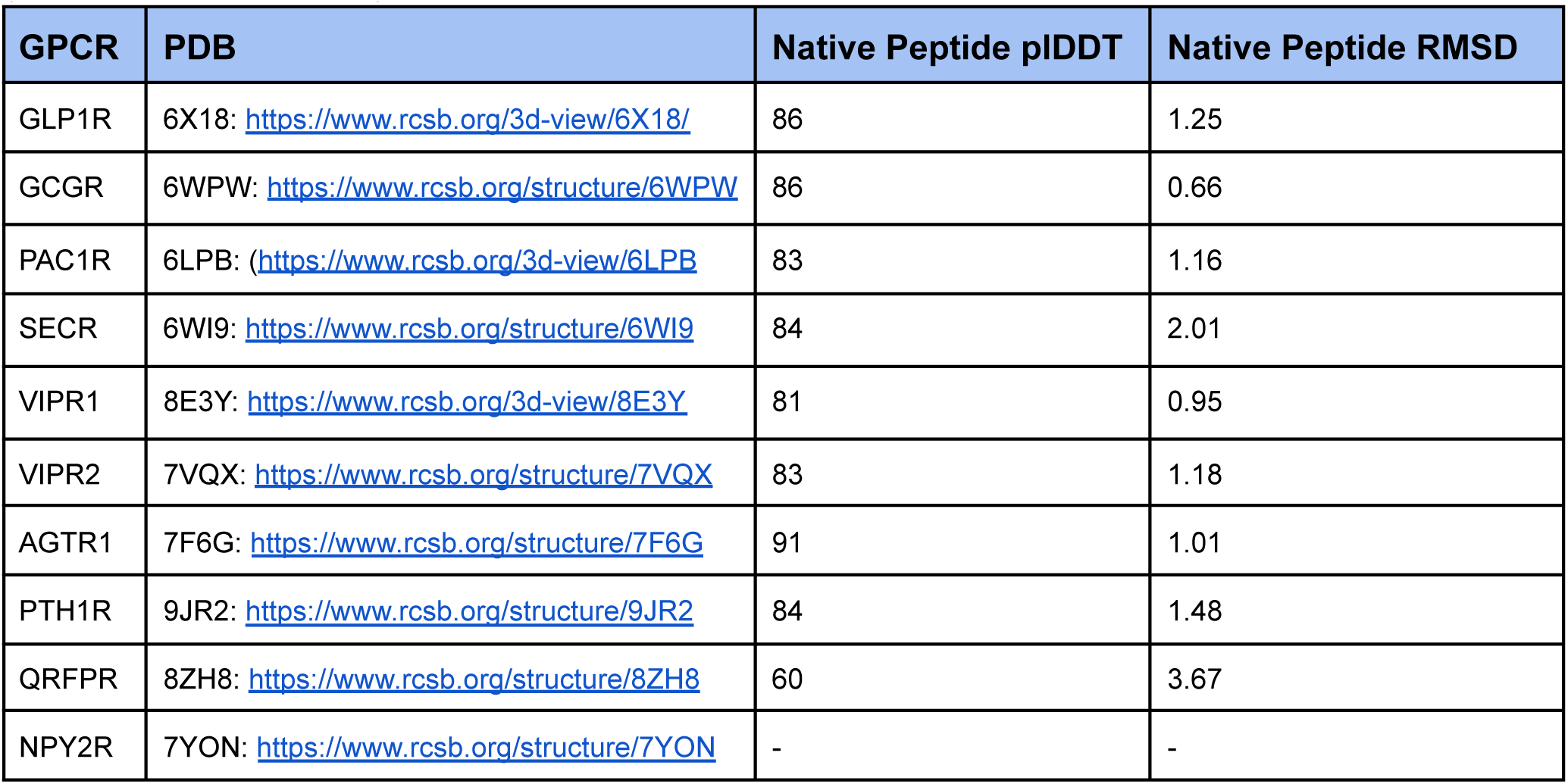
Selected GPCRs-peptide complexes (n=10) and their PDB IDs that can be accurately predicted with RareFoldGPCR, and one (8ZH8) that is not accurately predicted, and one that was not scored due to containing modified amino acids not available in RFG (7YON, amino acid TYC).

For each complex, we initiated binder design from a random sequence and applied mutation steps following the EvoBind protocol [17] (Figure 2a). We performed 10 independent design trajectories per target, each consisting of 1000 mutation steps (n = 10’000 sequences per GPCR). Figure 2b shows the evolution of the loss function (equation 1), indicating the emergence of structural features characteristic of functional binders as the loss decreases.

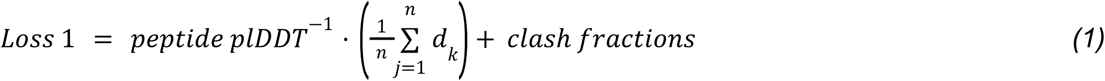

**Figure 2.**
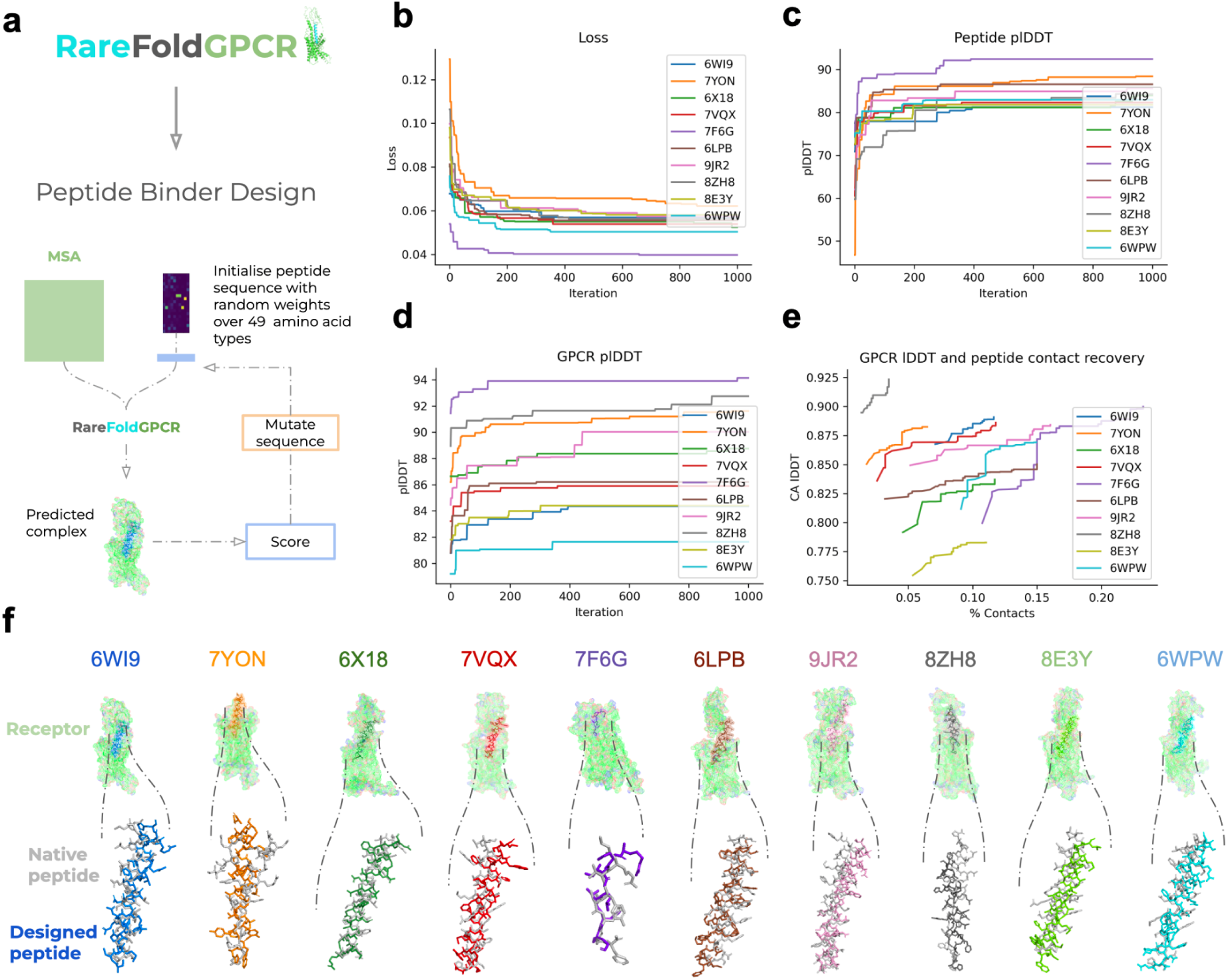
Peptide binder design with RareFoldGPCR. **a)** Overview of the peptide design pipeline using RareFoldGPCR. Starting from a multiple sequence alignment (MSA), peptide sequences are initialised randomly and iteratively mutated to optimise predicted binding to a GPCR, with candidate complexes scored using RareFoldGPCR. **b)** The loss function (equation 1) decreases over iterations for all targets, showing optimisation convergence. **c)** Best peptide plDDT (predicted local distance difference test) scores increase over the course of design iterations, indicating improved structural confidence. **d)** GPCR plDDT scores also improve during peptide design, suggesting that better peptide structures stabilise the receptor conformation, which remains relatively static. **e)** Correlation between GPCR Cα-lDDT (predicted similarity to the active state) and the fraction of native contacts recovered in the designed peptide. The similarity increases with the fraction of recovered contacts, indicating that agonistic modes are recognised. **f)** Examples of designed peptide-GPCR complexes for ten different targets. The receptor (green), native peptide (grey), and designed peptide (coloured) are shown. PDB IDs are indicated above each complex. The NPY2R complex (PDB: 7YON) shows a distinct binding mode, likely due to the presence of unusual modified amino acids in the peptide binder found in the PDB.

Here, the peptide plDDT refers to the average predicted local distance difference test score across all residues in the peptide, providing a measure of local structural confidence, with higher plDDT values indicating greater confidence in the predicted local geometry. The value d□ denotes the shortest distance between any atom in peptide residue k and any atom in the target protein, reflecting proximity to the receptor. Clash fractions measure the proportion of atoms in steric conflict, defined as interatomic distances below 1.5 Å for inter-chain contacts and below 1 Å for intra-chain contacts. Notably, all designs achieve plDDT scores above 80, regardless of prediction accuracy **(Table 1)**, indicating that the model can generate structurally confident predictions and discover novel solutions it deems plausible.

Peptide plDDT scores increase throughout the optimisation (Figure 2c) and, interestingly, the plDDT of the GPCR also rises during design (Figure 2d). This indicates that as the peptide adopts a more plausible conformation, the predicted GPCR structure becomes more stabilised. The trend is further supported by the increase in Cα lDDT relative to the known active state structure as contact recovery improves, suggesting that the network has learned features that drive specific receptor states (Figure 2e). However, contact recovery remains at only 20 per cent, indicating that the exact native interactions are not reproduced. Instead, the network generates novel interactions that are nonetheless assigned high confidence.

Together, these trends suggest that high peptide plDDT scores are predictive of both native-like contacts and accurate receptor states. In some cases, however, contact recovery remains low. Such cases may represent alternative or novel binding modes, as previously observed with EvoBind [18], where successful binders diverged from native contacts while still achieving functional outcomes. It is also possible that accurate modelling of the receptor’s active state is a prerequisite for recovering the specific interactions required for activation. Notably, the lowest loss is obtained for 7F6G (see Figure 2f for predicted structures), which also shows the highest contact recovery and peptide plDDT. This points to receptor activation being, at least in part, a search problem. We next test this experimentally using GLP1R as a model system.

### Peptide Agonist Design with NCAAs

To more carefully evaluate the design capabilities of RFG, we explore the possibility of designing novel agonists that include NCAAs towards GPCRs and validate these experimentally. If successful, this would indicate that entirely novel interfaces can be generated and GPCRs activated in ways beyond what is known from nature through RFG. We selected the native glucagon-like peptide-1 receptor (GLP1R) in complex with its endogenous ligand GLP-1 (PDB ID: 6X18) as a model system for peptide agonist design. GLP1R plays a central role in metabolic diseases such as type 2 diabetes and obesity [1] and belongs to class B2 GPCRs, a therapeutically important receptor family. We designed both linear and cyclic agonists for GLP1R by partially scaffolding residues from the native GLP-1 peptide and using a combination of canonical and NCAAs to complete the remaining sequence (Figure 3).

**Figure 3.**
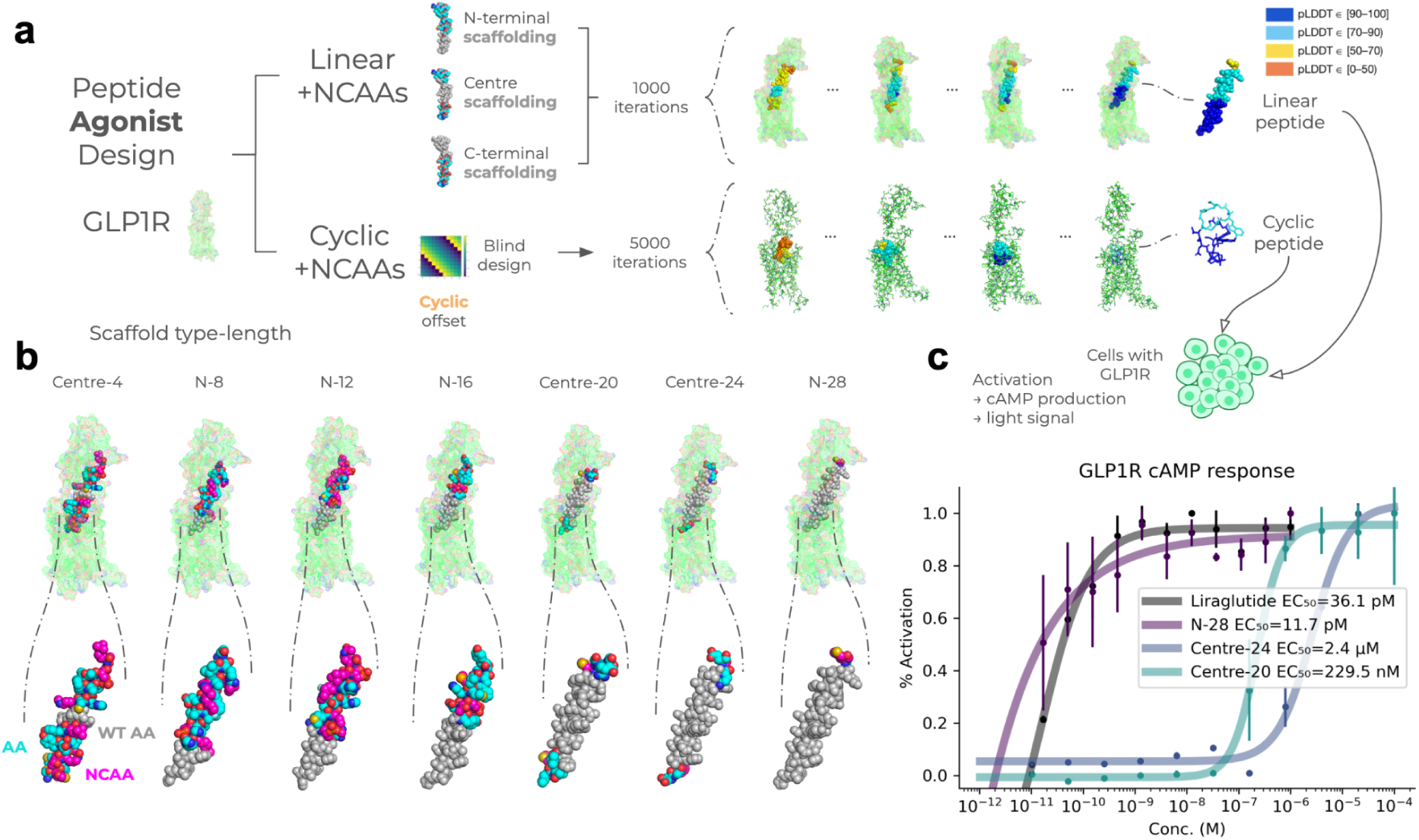
Linear and Cyclic agonist design for GLP1R. **a)** Overview of the design strategy. For scaffolding, native GLP-1 residues are retained in different regions (N-terminal, central, or C-terminal scaffolds) while the remaining sequence is designed using canonical and NCAAs. For the cyclic case, no scaffolding is used. The procedure is entirely blind, and any combination of amino acids in any cyclic conformation is designed to obtain a low design loss (Methods). **b)** Visualisation of selected linear scaffold designs, showing full GLP1R complexes to highlight diverse binding modes incorporating both regular and NCAAs. Designed peptide regions are shown in magenta for noncanonical residues and cyan for canonical ones, with scaffolded residues in grey. Scaffold lengths 4 and 8 could not be easily synthesised. **c)** Functional validation of selected linear designs in GLP1R cAMP cell assay (see Supplementary Figure 8 for all validated designs). All assay curves are shown in overlay alongside the reference agonist liraglutide (Lira). The best agonist is N-28, slightly outperforming Lira (EC₅₀ of 16.9 vs 36.2 pM). Interestingly, both Centre-20 and Centre-24 show potent agonist activities (229 nM and 2.4 μM, respectively,) showing that functional outcomes are obtained even without scaffolding the important N-terminal. This is important as the enzyme DDP-4 cleaves the first two residues of GLP-1, leading to rapid turnover, but this can be avoided with centre scaffolding. The lines represent sigmoidal fits, and the points represent averages from three replicates with vertical lines as standards.

For the linear designs, we systematically explored scaffolding strategies by selecting continuous stretches of 4 to 28 residues (with a step size of 4) from either the N-terminus, the central region, or the C-terminus of GLP-1. The remainder of the sequence (to a total length of 30 residues, matching the full GLP-1 in PDB ID 6X18) was filled through sequence design using both canonical and NCAAs (Methods). This allows us to evaluate the precise design capabilities of expanding on natural hormones with NCAAs in different ways. We find that high-confidence designs are generated in all cases, although the N-terminal and centre scaffolding yield lower losses than the C-terminal (Supplementary Figure 3). This is related to the interaction with the orthosteric pocket, which begins at the N-terminal, making the C-terminal harder to scaffold and the RMSD of the scaffold region thereby higher (Supplementary Figure 4).

In the cyclic case, the design problem becomes more challenging due to the conformational constraints imposed by cyclisation. To explore this case, we scaffold 2, 4, 6, 8, 10, 12, or 14 residues from the native GLP-1 peptide and use a total peptide length of crop size × 2 + 4 (i.e., 8, 12, 16, 20, 24, 28, 32 residues). However, we find that scaffolding cyclic peptides in this way performs poorly, likely because the original binding mode of the linear peptide cannot be maintained after cyclisation (Supplementary Figure 5). Fortunately, the sequence-based nature of the design process in RFG enables the discovery of entirely new binding modes [18]. Therefore, we performed an untargeted de novo design of cyclic peptides with lengths ranging from 15 to 30 residues, running 5000 iterations. This approach successfully generated high-confidence binders (Supplementary Figure 6), demonstrating that RFG can design novel cyclic peptide agonists beyond the constraints of the native scaffold (see the next section for the cyclic selection and experimental results).

For the linear peptide design, we selected sequences with a plDDT > 80 and the best design loss per scaffold length (n = 7, Tables 2 and 3, Figure 3b) for synthesis and subsequent experimental validation in GLP1R cAMP activation assays (Methods). This selection ensures high-confidence designs while covering diverse scaffold lengths for comprehensive testing. However, scaffold lengths 4 and 8 could not be easily synthesised, and these designs are not included in the experimental validation. Both N-terminal and centre scaffolding display activation, and the designs with longer scaffolded portions display lower EC₅₀ values (as expected, Figure 3c). However, scaffolding a larger part does not guarantee a lower EC₅₀, as centre-24 reports an EC₅₀ of 2.4 μM and centre-20, 229 nM. This highlights the difficulty of the problem as a single residue substitution can disrupt the agonist activity.

**Table 2.**
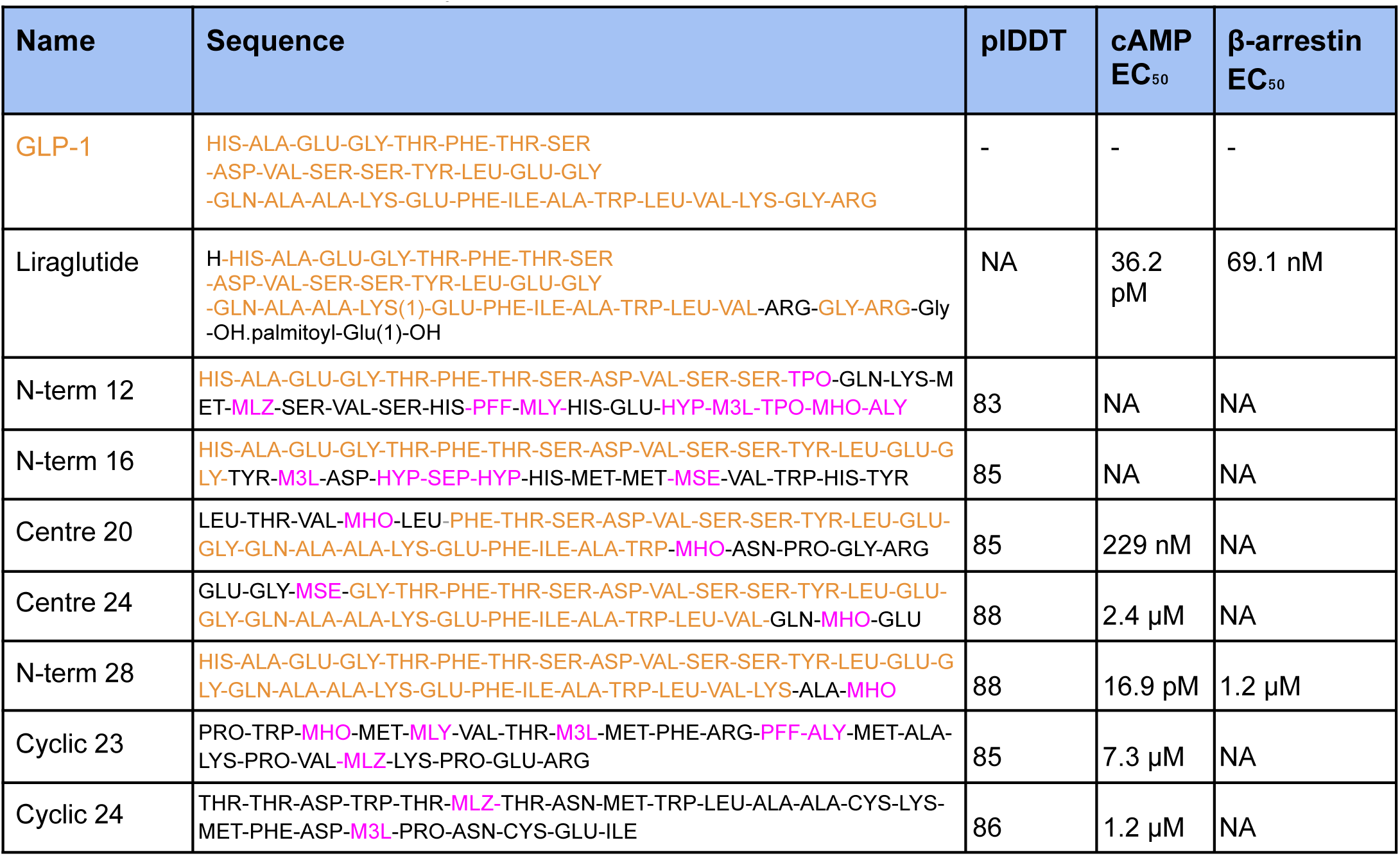
GLP1R cAMP/β-arrestin EC₅₀ values. The scaffolded part of the natural hormone (GLP-1) is highlighted in orange. The commercial Liraglutide has scaffolded almost the entire GLP-1 sequence, but introduced modifications (e.g. to LYS(1)). The native GLP-1 was not evaluated here, indicated by “-”.

**Table 3.**
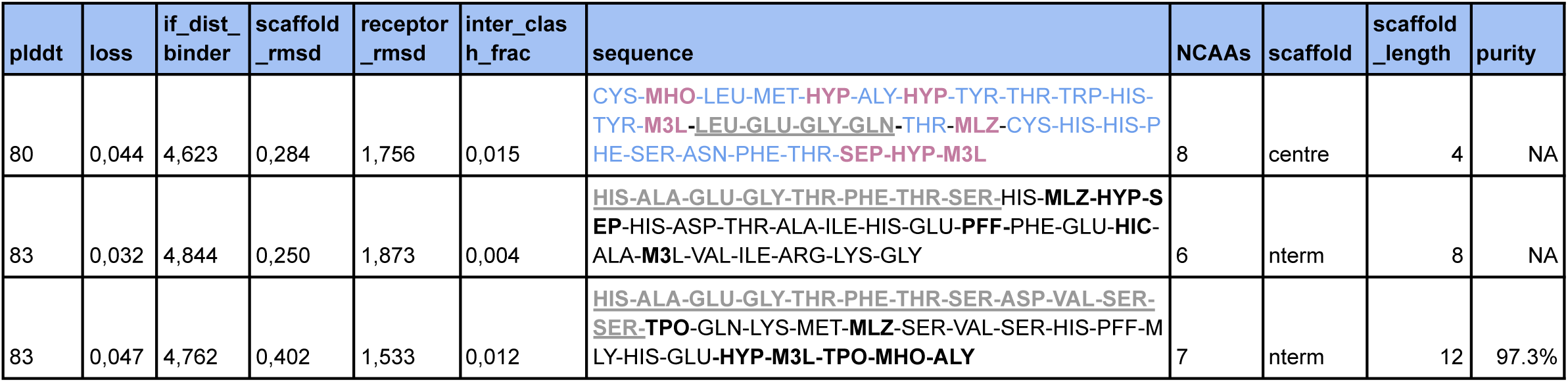
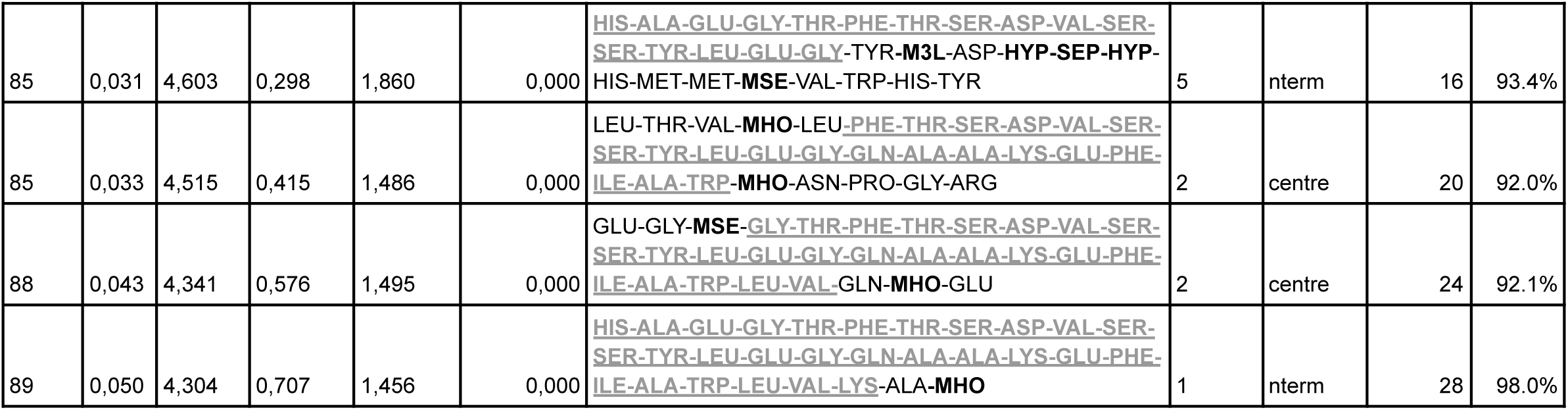
Linear selection. Summary of scaffold-based linear peptide designs after filtering for plDDT > 80 and loss improvement. Intra-chain clash fractions were excluded from the table as they were consistently low. Complete metrics for all runs are provided in the Data Availability section. Scaffold lengths 4 and 8 could not be easily synthesised.

**Table 4.**
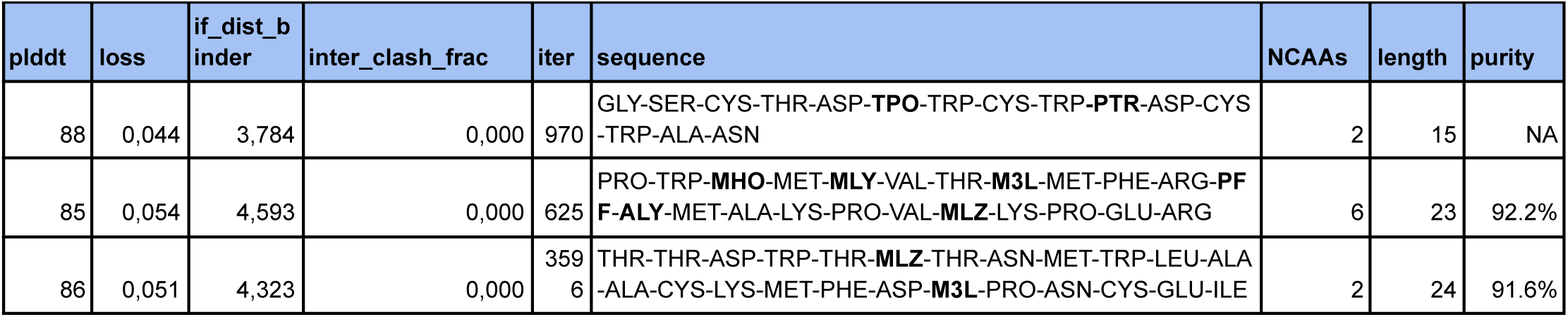
Cyclic selection. Summary of cyclic peptide designs after filtering for plDDT > 80 and loss improvement. Intra-chain clash fractions were excluded from the table as they were consistently low. Complete metrics for all runs are provided in the Data Availability section. Length 15 could not be easily synthesised.

The results show that we can scaffold different parts of natural hormones like GLP-1, introduce unseen NCAA into the interface and obtain functional outcomes. The successful centre scaffolding suggests that function can be retained without scaffolding the important N-terminal and thereby avoiding N-terminal degradation from DDP-4 [26,27], which renders the natural GLP-1 inactive by cleaving off only 2 residues (HA-). The best EC₅₀, observed for

N-28, outperforms the commercial Liraglutide slightly (16.9 vs 36.2 pM), and all but N-12 and N-16 display agonist activities with EC₅₀ values in the μM-pM range (see Supplementary Figure 8 for all validated linear designs). The fact that both N-12 and N-16 fail shows that even if the part of the native peptide that engages the orthosteric pocket is scaffolded, there is no guarantee of getting an agonist. Agonist design is still very difficult. Next, we analyse the agonist spectrum to the extreme by designing entirely new sequences, topologies and activation modes through blind cyclic agonist design.

### Cyclic Agonist Design with NCAAs

The scaffolding approaches used for the linear design could not accommodate the conformational constraints required for cyclic topologies. The binding mode of the linear GLP-1 agonist is incompatible with cyclisation in the structural modelling, likely necessitating both a new binding mode and potentially a different mechanism of receptor activation. As a result, successful cyclic peptide agonist design requires *de novo* design without relying on native residue scaffolds.

As previously demonstrated with the standard 20 amino acids [18], novel binding modes can emerge that are capable of activating GPCRs. Therefore, we performed untargeted cyclic peptide design across lengths of 15-30 residues and selected the best designs per length.

This approach investigates the possibility of designing agonists without prior knowledge of either agonist lengths or topologies. Applying a plDDT > 85 cutoff for structural confidence reduced the set to two cyclic peptide lengths that were synthesised successfully (23 and 24 residues) from which we selected the designs with the lowest losses according to equation 1 (Figure 4a).

**Figure 4.**
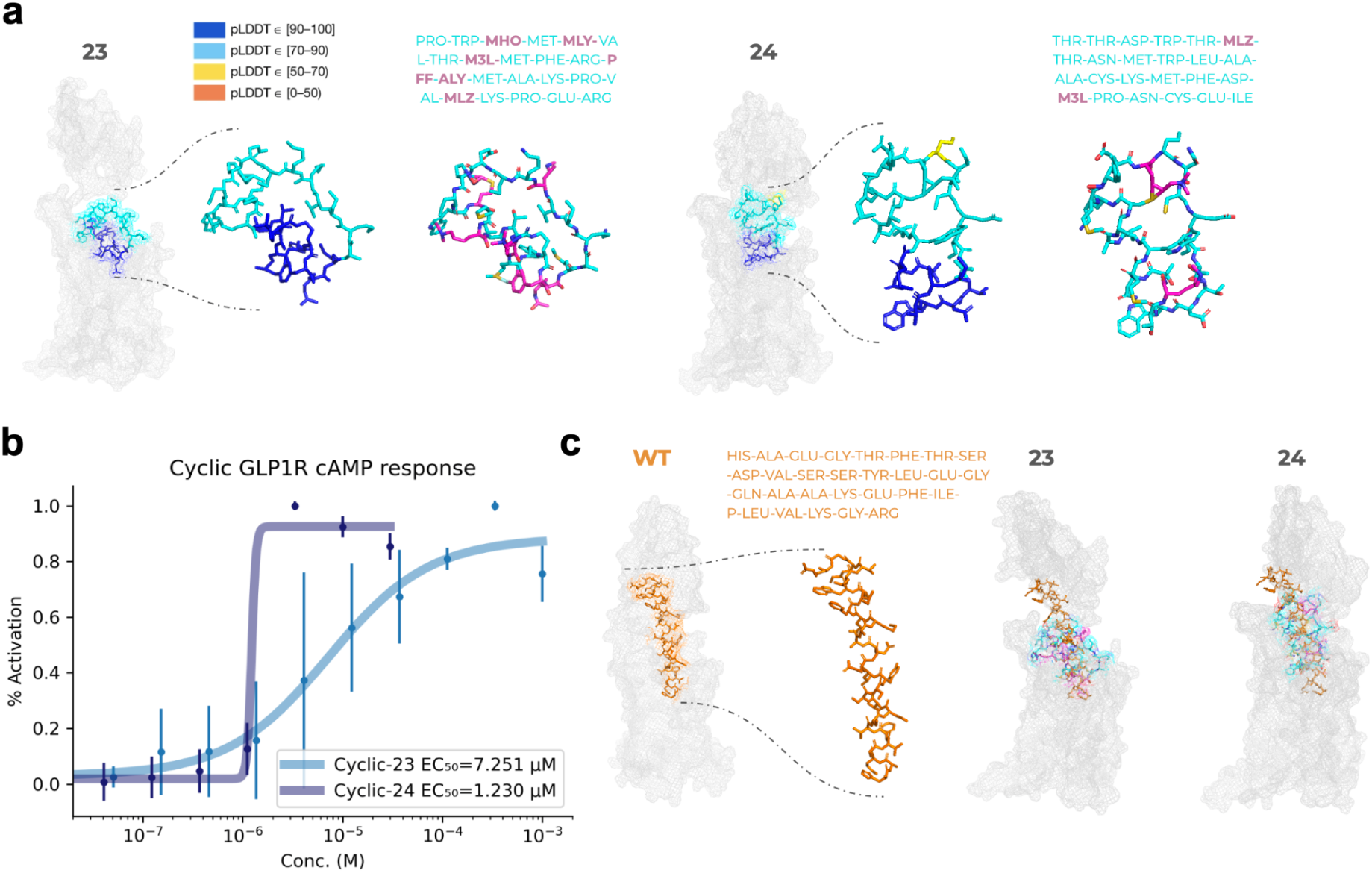
Experimental results and predicted structures of the cyclic agonist designs for GLP1R. **a)** Cyclic peptide designs selected for synthesis. Predicted complexes are shown with the receptor in mesh representation and the designed peptide in stick format, coloured either by plDDT (confidence) or by amino acid type (NCAAs in magenta, canonical residues in cyan). Notably, all NCAAs contribute directly to interactions within the orthosteric pocket. The plDDT values are also highest down into the orthosteric site. **b)** Cell-based cAMP assay results showing that both cyclic peptides act as potent agonists with activities in the low μM range. Cyclic-23 has an EC₅₀ of 7.3 μM, and Cyclic-24 1.2 μM. The raw data, relative light units, are visualised in Supplementary Figure 9. The lines represent sigmoidal fits, and the points represent averages from three replicates with vertical lines as standards. **c)** Structural comparison of the predicted cyclic designs with the wild-type GLP-1 peptide (orange, experimental structure PDB ID: 6X18). The cyclic peptides adopt binding modes that are distinct from each other and from GLP-1, demonstrating that RFG enables the discovery of novel chemistries and activation mechanisms. This also suggests that much of the agonist space may be unexplored and that transfer learning can generalise to novel activation.

Experimental validation in GLP1R cAMP activation assays shows that both length 23 and 24 activate GLP1R with EC₅₀ values of 7.3 and 1.2 μM, respectively (Figure 4b, Table 2). The sequence overlap with the WT GLP-1 is missing, and the topology of the designs is novel, with NCAA incorporated into the predicted interfaces (Figure 4c). The designs themselves are also different from each other in both sequence and structure, indicating that RFG can generalise to create multiple novel agonistic modes.

The CLUSTAL O(1.2.4) [28] multiple sequence alignment (with X denoting NCAAs and matches shown in bold) reveals that only a few residues from the linear WT sequence align with the cyclic designs, and large gaps are required to accommodate these limited matches. Moreover, the difference in linear versus cyclic topologies implies that such sequence-level similarities would not translate into equivalent 3D arrangements, consistent with the distinct binding modes predicted (Figure 4c). Together, these observations suggest that a substantial fraction of GPCR agonist space remains unexplored, and that transfer learning enables generalisation to entirely novel activation modes with NCAAs.

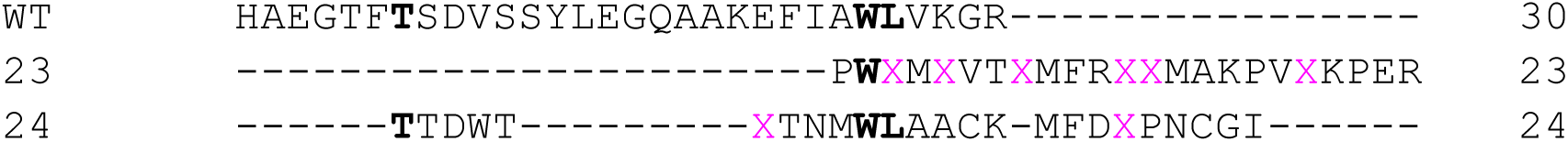

### Pathway specificity: cAMP and β-arrestin

When an agonist binds to GLP1R, it triggers two key signalling pathways: the G-protein pathway, which leads to the creation of cAMP, and the β-arrestin pathway, which helps with receptor desensitisation and other independent signalling. The idea of biased agonism, where a ligand mainly activates one pathway over the other, is important in drug development [29] and may help separate the desired therapeutic effects from unwanted side effects [30]. To investigate the structural basis for pathway bias, we analysed metrics related to the RFG-predicted peptide-receptor complex structures.

We find an interesting relationship between GLP1R β-arrestin and cAMP activation, where peptides with pM cAMP EC₅₀ that scaffold more of the WT (N-28) are likely to activate both pathways, while the others only induce the cAMP pathway (Figure 5a, Table 2). Interestingly, N-28 activates the cAMP pathway more strongly than Liraglutide while having a weaker activity at the β-arrestin pathway, suggesting a more favourable signalling profile (Table 2).

**Figure 5.**
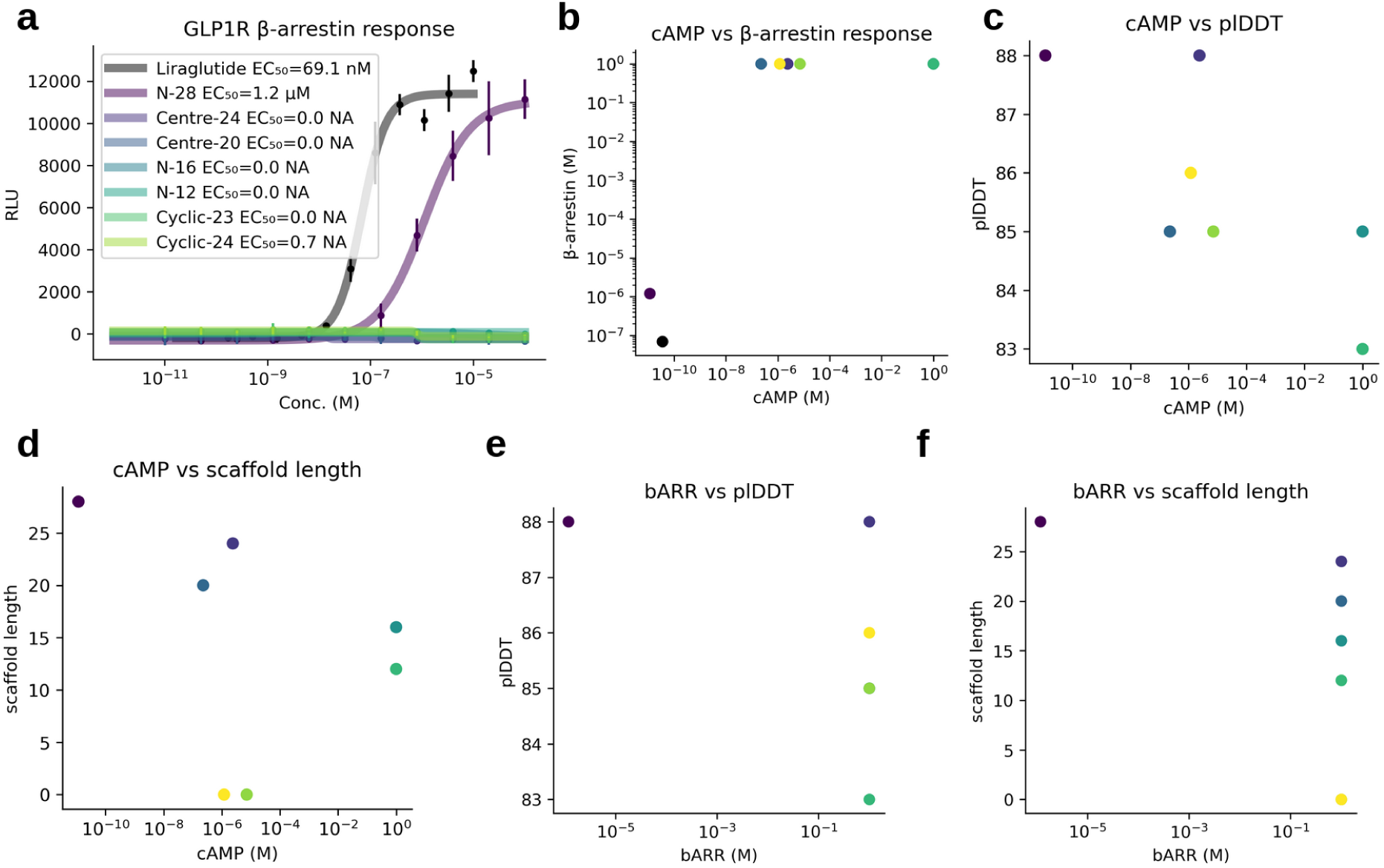
GLP1R cAMP and β-arrestin activation. **a)** β-arrestin response curves for designed peptides and reference ligands (RLU=Relative Light Units vs concentration). Only N-28 and Liraglutide show measurable activation. Liraglutide activates with an EC₅₀ of 69.1 nM, compared to 1.2 µM for N-28, despite N-28 being more potent in the cAMP pathway, indicating a more favourable signalling profile. The lines represent sigmoidal fits, and the points represent averages from three replicates with vertical lines as standards. **b)** Comparison of cAMP and β-arrestin EC₅₀ values for all tested peptides. Most designs show strong cAMP activity but lack β-arrestin recruitment, consistent with pathway bias. Missing cAMP/ β-arrestin activation is set to 1. **c)** Relationship between peptide plDDT and cAMP EC₅₀. Higher plDDT values generally correspond to lower EC₅₀, indicating that structural confidence correlates with functional potency. Missing cAMP activation is set to 1. **d)** Scaffold length vs cAMP EC₅₀. Longer scaffolds tend to achieve higher potency in the cAMP pathway, while cyclic peptides (assigned scaffold length = 0) cluster in the low µM range. Missing cAMP activation is set to 1. **e)** Relationship between plDDT and β-arrestin EC₅₀. Only N-28 displays measurable values, limiting correlation but suggesting a requirement for extended scaffolds. Missing β-arrestin activation is set to 1. **f)** Scaffold length vs β-arrestin EC₅₀. Only N-28 is active, supporting the observation that longer scaffolds more closely resembling the native ligand are necessary for β-arrestin recruitment. Missing β-arrestin activation is set to 1.

**Figure 6.**
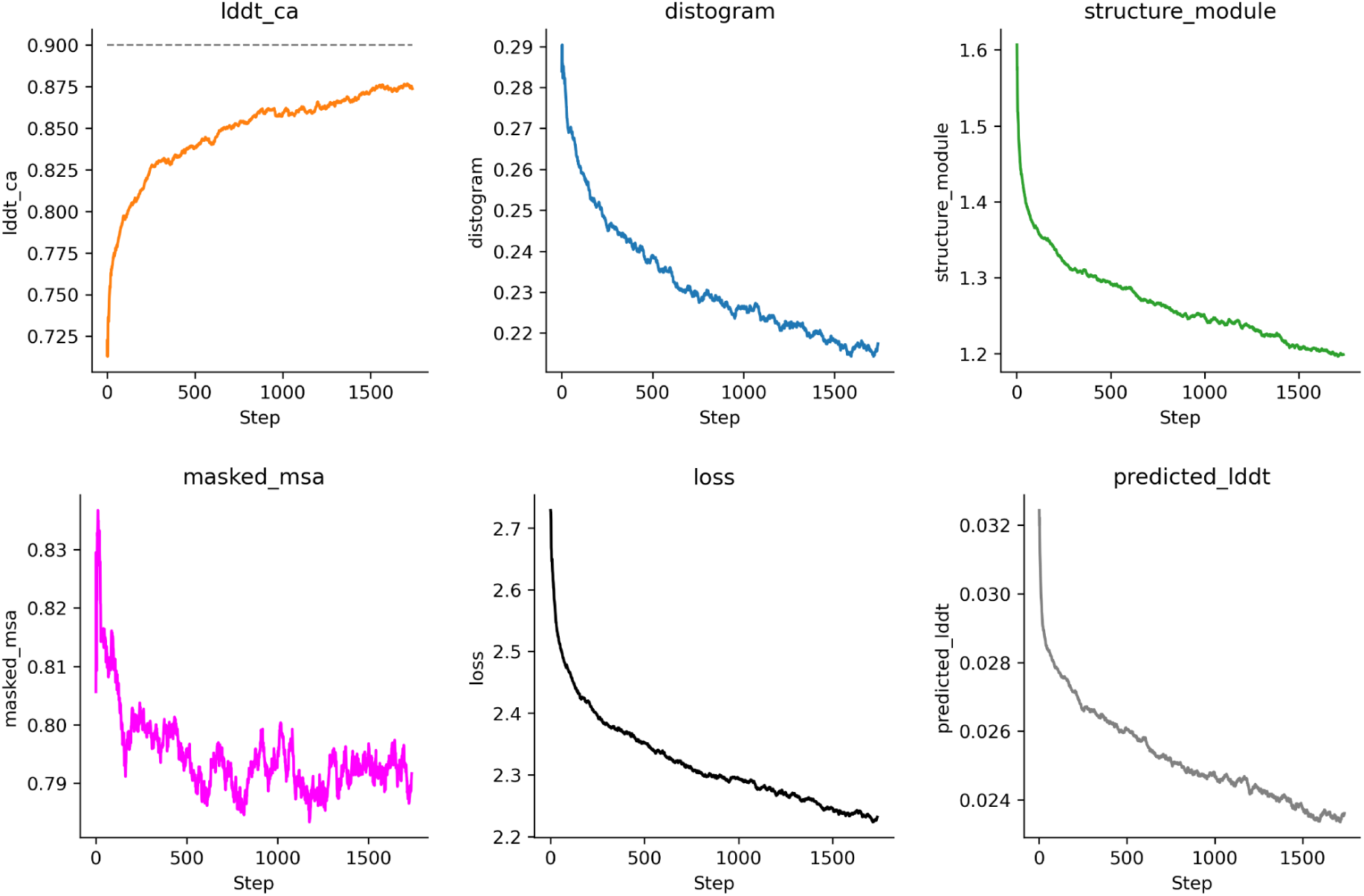
Training curves for GPCR tuning. Training losses and metrics are shown with a smoothing window of 50. Loss components include the structure_module (AUX, FAPE and angular terms); distogram (pairwise distance prediction), combined loss (equation 1), and predicted_lddt (local distance confidence), following AlphaFold2 [14].

**Figure 7.**
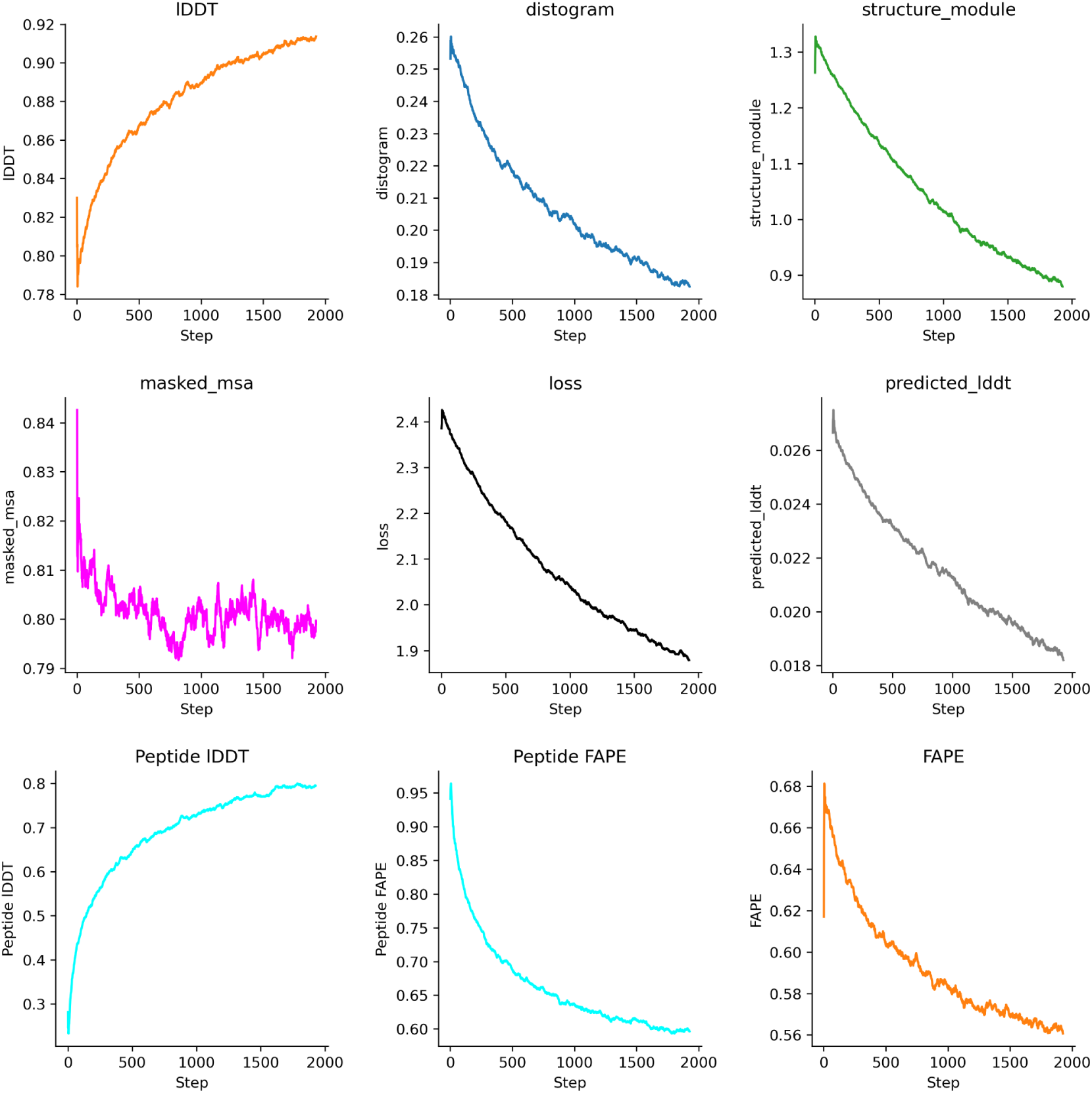
Training curves for GPCR-peptide complex tuning. Training losses and metrics are shown with a smoothing window of 50. Loss components include the structure_module (AUX, FAPE and angular terms); distogram (pairwise distance prediction), combined loss (equation 1), and predicted_lddt (local distance confidence), following AlphaFold2 [14]. In addition, we show the peptide lDDT and FAPE as well as the FAPE over the whole complex. Comparing the peptide-specific scores with those over the entire peptide-GPCR complex shows that it is more difficult to learn the peptides.

Peptides with shorter scaffolds, N-12 and N-16, completely lost their agonist function in both cellular assays despite incorporating WT parts known to interact with the orthosteric pocket (Figure 3b).

When RFG is allowed to generate agonists blindly, as in the cyclic case, potent low μM cAMP EC₅₀s are obtained, but no β-arrestin recruitment. This suggests that RFG has internalised cAMP pathway specificity for GLP1R and also outlines a complex agonistic relationship where scaffolding larger WT portions may be necessary to obtain full agonistic profiles across all downstream pathways. Compared to the linear WT, the cyclic designs are predicted to significantly expand the orthosteric pocket (Figure 4c). In addition, no engagement with the extracellular (EC) domain is predicted to happen. We hypothesise that engaging the entire orthosteric pocket and EC-domain comparable to the WT is necessary for the full signalling profile, as the centre scaffolds also don’t engage the β-arrestin pathway, and the plDDT being lower in the linear non-scaffolded regions (Figure 8).

**Figure 8.**
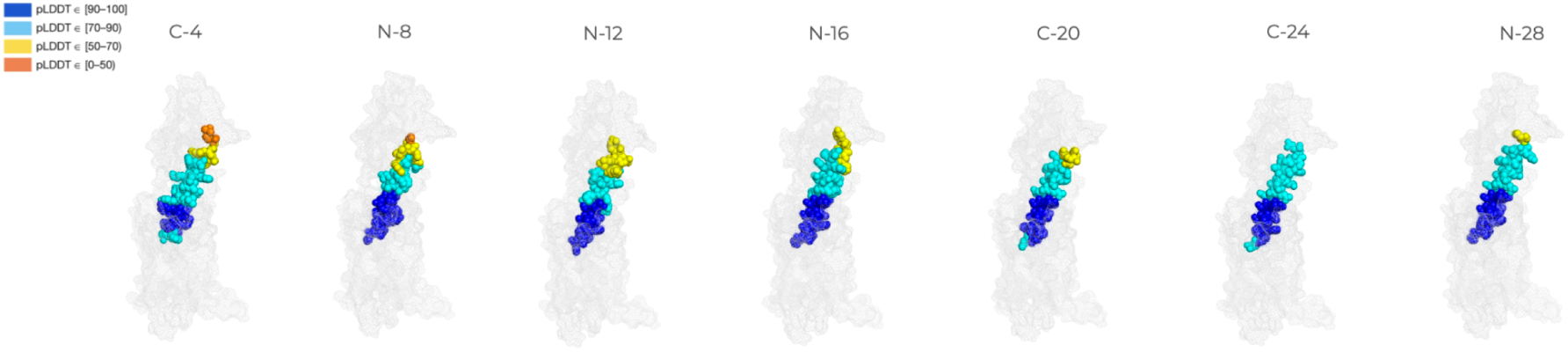
Linear selection of peptides for synthesis coloured by plDDT. The scaffold length and type (C=Centre, N=N-terminal)

Analysis of design metrics further supports these trends (Figure 5c-f): longer scaffolds generally yield lower EC₅₀ values and higher plDDT scores, consistent with more native-like binding, yet only N-28 elicits measurable β-arrestin recruitment. This highlights both the difficulty of designing full agonists across pathways and the potential of RFG to bias signalling toward desired outputs.

## Discussion

RareFoldGPCR (RFG) demonstrates the power of transfer learning: despite no training data for GPCR agonists with NCAAs and only single-chain inputs, the model accurately places rare chemical groups into agonist-binding pockets. Here, we use GLP1R as a proof-of-concept to investigate the specific potential of RFG in various scenarios with *in vitro* validation. By varying scaffold length and region in GLP-1, we probe RFG’s ability to modify known agonists and, with cyclic topologies, generate entirely novel agonists. This reveals how RFG internalises GPCR activation principles and enables both expansion of natural agonists and *de novo* agonist design.

For linear designs, 1000 optimisation iterations were sufficient to recover active agonists, whereas 5000 were required for cyclic peptides. Additional iterations consistently increased peptide plDDT and may further improve functional outcomes. As expected, longer scaffolds closer to the native ligand yielded stronger cAMP activation, reflecting the challenge of outperforming nature. Interestingly, scaffolding N-terminal regions of GLP-1 that engage the orthosteric pocket (e.g. N-12 and N-16) did not consistently yield agonists, underscoring both the difficulty of designing novel activators and the exquisite sensitivity of these interactions. Nevertheless, the use of NCAAs enables the introduction of novel chemistry and cyclic topologies, pointing towards alternative activation modes and potential signalling bias.

For GLP1R, our results highlight the potential to probe β-arrestin recruitment, receptor desensitisation, and pathway-selective agonism. Importantly, even peptides with weaker cAMP activation can be valuable if they show reduced β-arrestin recruitment, as this may mitigate desensitisation and sustain signalling. This is exemplified by the cyclic RFG designs, which consistently activate cAMP at low micromolar potency while showing no β-arrestin activity, pointing to a pathway-selective mechanism that could be therapeutically advantageous. Conversely, N-28, which scaffolds more of the WT GLP-1 sequence, activates both cAMP and β-arrestin pathways, with stronger cAMP activity than Liraglutide yet weaker β-arrestin signalling.

We hypothesise that engaging the entire orthosteric pocket and EC domain, as in the WT, is required for achieving the full signalling profile. In contrast, designs incorporating NCAAs that introduce localised perturbations near the signalling interface may bias the receptor toward the Gα-coupling conformation, explaining the absence of β-arrestin recruitment. These findings suggest that RFG can enable mechanistically guided exploration of biased agonism for GLP1R, supporting rational tuning of both potency and pathway specificity through NCAAs. This offers a path to ligands that prioritise durable efficacy over maximal binding strength, though validation *in vivo* will be essential.

Beyond activity, NCAAs can also confer proteolytic resistance (e.g. avoiding DPP-4 cleavage [26,27]) and reduced immunogenicity, as suggested by earlier RareFold studies [19] and general findings [31,32]. While these aspects remain to be validated *in vivo* in future studies, RFG already enables fully de novo cyclic agonist design, pointing toward stable, long-acting, and potentially immunogenically silent therapeutics. The RFG approach is also general and scalable as it supports efficient parallel runs (> 10 designs on a single A100 GPU), and in silico analysis (Figure 2) indicates broad generalisation across GPCR families. Though GLP1R served as a test case, RFG has the potential to extend to any receptor, including orphans, since it is purely sequence-based. This provides a powerful route to systematically explore agonist sequence space beyond natural amino acid chemistry, enabling rational design of next-generation GPCR therapeutics.

## Methods

### Data from GPCRdb

We retrieved all available GPCR structures from GPCRdb (https://gpcrdb.org/structure/, accessed 11 March 2025), obtaining 1650 entries. Of these, 1645 structures were successfully downloaded in .cif format from the PDB, and 1634 could be parsed successfully with BioPython [33] (99.3% recovery). All structures contained at least 200 residues (Supplementary Figure 7).

To reduce redundancy and ensure representation of small structural variations, we clustered the sequences at 80% sequence identity using MMseqs2 [34] with the command:

~~~
mmseqs easy-cluster examples/DB.fasta clusterRes tmp --min-seq-id0.8 -c 0.8 --cov-mode 1
~~~

This yielded 316 sequence clusters, which were used to sample representative structures during training.

We further retrieved all GPCR-peptide complexes from GPCRdb (accessed 13 March 2025), identifying 572 receptors bound to peptide ligands. For each complex, the peptide chain shorter than 50 amino acids with the most receptor contacts was extracted, yielding 315 valid complexes after excluding one unscorable entry due to only containing modified unusual residues (PDB 8XIQ). The corresponding receptors clustered into 83 groups at 80% sequence identity.

### Fine-tuning RareFold for GPCRs

The RareFold model [19] was first fine-tuned on GPCR structures using the receptor clusters sampled according to the inverse cluster size, with a batch size of 16, crop size of 384 and learning rate of 10⁻⁴. We used the same losses as in AlphaFold2 [14] and RareFold [19] (equation 2, Figure 6). The best validation loss on the orthogonal problem of GPCR-peptide complex prediction was obtained after 500 steps (Figure 1).

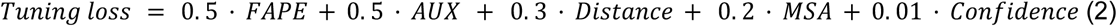

Where *FAPE* is the frame aligned point error, *AUX* a combination of the *FAPE* and angular losses, *Distance* a pairwise distance loss, *MSA* a loss over predicting masked out MSA positions and *Confidence* the difference between true and predicted lDDT scores. These losses are defined exactly as in AlphaFold2 and we refer to the description there [14].

The weights from step 500 were then further fine-tuned on the peptide-bound subset to specialise the model for peptide-GPCR interactions. The same conditions and sampling as applied for the single chain model were used with the difference of introducing an additional chain. We enabled this possibility by treating the second chain as a disconnected part of the same protein using a “chain gap”. This is possible as the maximal relative positional encoding is 32, which means that any jump in the consecutive residue numbering larger than 32 will result in the model interpreting a chain gap, which here means the free peptide chain. In this way, we enable transfer learning of a single chain model to the complex problem of GPCR-peptide structure prediction.

Figure 7 displays the losses used for the complex fine-tuning with the additional peptide lDDT and FAPE, highlighting the difference between the overall and peptide learning outcomes. The peptide FAPE is higher than that of the whole model, indicating that the more flexible and less structured peptides are more difficult to learn and place in relation to the GPCRs. Another factor is that the peptides don’t have coevolutionary information to represent them. Throughout this work, all peptides are represented as single sequences.

This also enables RareFoldGPCR to learn how single-sequence peptides may interact with the target receptors without relying on coevolution - essential for accurate prediction and design in novel settings.

### Scoring GPCR-peptide interactions

Predictions were run with three recycles throughout. To account for large chain breaks commonly observed in GPCR structures (e.g. GLP1R) due to disordered extracellular regions, we used the structural residue index with gaps included, ensuring continuity of residue numbering and avoiding artificial truncation between membrane and extracellular domains. Model quality was assessed using GPCR Cα-lDDT and peptide Cβ-RMSD, with glycine residues treated as Cα. For peptide evaluation, Cβ atoms within 8 Å of the receptor were aligned, and complexes with peptide Cβ-RMSD <2 Å were considered successful. The number of successful complexes increased with tuning, yielding 71, 99, 126, and 145 at steps 200, 500, 1000, and 1500, respectively. Parameters from step 1000 were selected for downstream peptide design.

### Mutation analysis

Peptides with accurate predictions (Cβ RMSD < 2 Å relative to the native ligand) were selected from the tuning runs, yielding 71, 99, and 126 peptides at steps 25700, 26000, and 26500, respectively. For each peptide, mutations were introduced systematically by generating 10 independent samples for each mutation number (e.g. 10 with one mutation, 10 with two, etc.). Structural evaluation was performed by computing peptide RMSDs based on Cβ atoms (Cα for glycine), consistent with the design evaluation. For peptides containing NCAAs not supported in RareFold, residues were mapped to *UNK*. When *UNK* was subsequently replaced by a standard amino acid, direct atom mapping could not be performed; in these cases, matching Cα atoms were used instead. This adjustment did not affect plDDT or global scoring.

### Blind Design for GPCRs with known peptide ligands

For blind design experiments, we selected GPCRs with known peptide ligands. Each design run was initialised with 10 random peptide sequences, with peptide lengths matched to the corresponding natural ligand. Optimisation was performed for 1000 iterations.

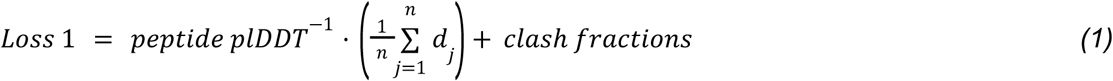

across all residues of the peptide [21], and dj is the shortest distance between any atom in peptide residue j and any atom in the receptor. The plDDT reflects local structural confidence, with higher values indicating greater reliability of the predicted conformation. Clash fractions quantify steric overlaps, penalising inter- and intramolecular atomic distances <1.5 Å and <1 Å, respectively.

### Contact recovery

To assess the similarity between designed and native peptides, we compared receptor–peptide contacts, defined as Cβ atoms (Cα for glycine) within 8 Å. Amino acids were grouped into five categories based on physicochemical properties (hydrophobic, polar, positive, negative, aromatic), such that different residues with similar characteristics were treated as equivalent (e.g. lysine and arginine both as positive). For each receptor interface position, the set of unique interaction categories observed in the native peptide was recorded. Contact similarity was then calculated as the fraction of these native interaction categories recovered in the designed peptide. Redundant interactions (e.g. a single receptor residue contacting two different hydrophobic residues) were only counted once, reflecting that maintaining one strong physicochemically relevant interaction may suffice to preserve binding.

#### Amino acid groupings

Hydrophobic: A, F, I, L, M, P, V, W, Y

Small: G

Polar: N, C, Q, S, T

Positive: R, H, K

Negative: D, E

### Linear and cyclic scaffolding design of GLP-1 with NCAAs

We implemented a scaffolding strategy where different regions of the native GLP-1 agonist (PDB ID: 6X18) were retained, while the remaining sequence was redesigned using a combination of canonical amino acids and NCAAs. This approach allows for controlled exploration of how much of the native binding information is required to maintain activity, while introducing new chemistry through noncanonical residues.

Specifically, we selected three types of scaffolds:

1. N-terminal scaffolds: retaining the first *x* residues of GLP-1.
2. Central scaffolds: retaining *x* residues from the middle of GLP-1.
3. C-terminal scaffolds: retaining the last *x* residues of GLP-1.

For the linear designs, lengths of the retained scaffold segments ranged from 4 to 28 residues in steps of 4 (i.e., 4, 8, 12, 16, 20, 24, 28 residues). For each scaffold length, the remaining positions up to the full 30-residue length of GLP-1 were designed de novo. This strategy allowed us to systematically investigate how the positioning and extent of the retained native sequence affect design performance. For each scaffold type and length, we generated 10 independent design trajectories starting from random initial sequences. We resample the MSA every 50 steps and run a total of 1000 mutation rounds (steps). We use the loss function:

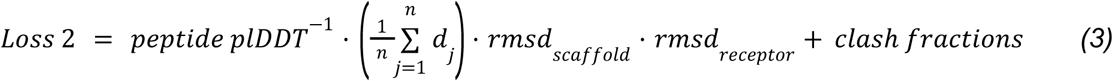

Here, the peptide plDDT is the average predicted local distance difference test score across the peptide residues, providing a measure of local structural confidence. The second term, represents the mean shortest distance between peptide atoms and any receptor atom, favouring close peptide-receptor contacts. The scaffold RMSD enforces structural fidelity by penalising deviations of the retained residues from their native Cα positions, while the receptor RMSD constrains the receptor backbone to its native state during design. Finally, the clash fractions penalise steric overlaps, defined as the fraction of inter-atomic distances below 1.5 Å (inter-chain) or 1.0 Å (intra-chain). Together, these terms balance structural confidence, receptor engagement, scaffold preservation, and physical plausibility.

For the cyclic peptide design, we explored the possibility of scaffolding 2, 4, 6, 8, 10, 12, or 14 residues from the native GLP-1 sequence while closing the peptide into a cycle. The total peptide length for each design was set to (scaffold size × 2) + 4, resulting in cyclic peptides of lengths 8, 12, 16, 20, 24, 28, and 32 residues. This approach aimed to preserve key native interaction motifs while allowing flexibility in the remaining sequence through the incorporation of canonical and NCAAs.

However, scaffolding proved unsuccessful in the cyclic case (Supplementary Figure 5). The structural constraints imposed by cyclisation appeared incompatible with the original linear binding mode of GLP-1, likely requiring not only new peptide conformations but also altered receptor interactions to achieve agonist activity. As a result, we shifted to a blind, fully de novo cyclic design without any scaffolding, following the loss function defined in Equation 1. This strategy allows the model to explore novel binding modes and receptor activation mechanisms that may not be accessible through scaffold-based approaches (see below).

### Linear selection

We selected sequences with plDDT > 80 and the lowest loss per scaffold length that showed improvement during optimisation. Out of 2926 designs with loss improvements, 1653 achieved plDDT > 80, and 1540 of these incorporated NCAAs. The most confident solutions were found when scaffolding was applied at the N-terminus (4 designs) or in the central region (3 designs), where the peptide extends into the orthosteric pocket (Figure 8).

Confidence also increased for longer scaffold lengths, likely reflecting the stabilising effect of retaining more native contacts in regions close to the binding site. The designs with the most NCAAs (scaffold lengths 4 and 8) could not be easily synthesised and are therefore excluded from the experimental validation.

### De novo cyclic peptide agonist design for GLP1R with NCAAs

To analyse the possibility of designing entirely novel binding modes, and as cyclic scaffolding proved unsuccessful (Supplementary Figure 5), we shifted to blind, fully de novo cyclic design without any scaffolding, following the loss function defined in *equation 1*. This strategy allows the model to explore novel binding modes and receptor activation. We designed binders of lengths 15-30 using 10 initialisations per length, and ran 5000 iterations with MSA resampling every 50 steps. Supplementary Figure 6 shows the optimisation curves.

### Cyclic selection

Cyclic peptide designs were generated with 5000 optimisation iterations and filtered using the same criteria as for the linear scaffolds (plDDT > 80 and loss improvement). For most designs, the top candidates were identified within the first 1000 iterations, although the best-performing case corresponded to a length of 24 residues. The peptide of length 15 could not be readily synthesised and was excluded from experimental validation.

### cAMP assay

To quantify cellular cAMP levels, the cAMP Hunter eXpress GPCR Kit (DiscoverX, 95-0062E2CP2L) was used. Cells were seeded into a 384-well plate (Corning, #3765) and incubated overnight. On the following day, cells were treated with either liraglutide as a control agonist or the designed cyclic peptides for 1 hour at 37°C, and the samples were further processed according to the manufacturer’s protocol. Briefly, following agonist incubation, 15 μL of cAMP Antibody Reagent was added to all wells of the assay plate. A cAMP Working Detection Solution was prepared in a separate 15 mL polypropylene tube by mixing 19 parts cAMP Lysis Buffer, 5 parts Substrate Reagent 1, 1 part Substrate Reagent 2, and 25 parts cAMP Solution D. Subsequently, 60 μL of the cAMP Working Detection Solution was added to all wells without pipetting up and down or vortexing the plates. The assay plate was incubated for 1 hour at room temperature in the dark to allow the immunocompetition reaction to occur. Next, 60 μL of cAMP Solution A was added to all wells without pipetting up and down or vortexing, followed by a 3-hour incubation at room temperature in the dark. Luminescence was measured using a standard luminescence plate reader with a read time of 1 second per well for the photomultiplier tube. All luminescence was normalised with the negative control values.

### β-arrestin assay

The β-arrestin recruitment assay was performed as previously described [18]. Briefly, HTLA cells were transfected in 100 mm dishes at a density of 6×10^6^ cells/dish with 20 µg of GLP1R plasmid DNA using the Calcium Phosphate Transfection Kit (Invitrogen, #K2780-01). Following transfection, cells were seeded at 15,000–20,000 cells per well in 40 µL of medium into poly-D-lysine-coated, 384-well, white, clear-bottom plates (Greiner Bio-one). Stock peptides were diluted to the desired concentrations in an assay buffer (HBSS containing 20 mM HEPES) and dispensed into the cell plates using a SELMA 384 liquid handler. After peptide incubation overnight, the peptide-containing medium (55 µL/well) was removed. Bright-Glo™ reagent, diluted 1:10 in the assay buffer, was then added (20 µL/well) using a Multidrop Combi dispenser at low speed. The plates were incubated for 20 minutes at room temperature in the dark before luminescence was measured for 1 second per well using a Hidex Sense microplate reader. All luminescence was normalised with the negative control values.

## Acknowledgements

PyMOL was used for structural visualisation.

## Contributions

P.B. designed the study, developed RareFoldGPCR, designed the binders, prepared the figures and wrote the manuscript. Q.L. performed the experimental validation with support from T.H and P.B.

## Data availability

All data and results used for this study are available at: https://zenodo.org/uploads/15180406

## Code availability

RareFoldGPCR is available at: https://github.com/patrickbryant1/RareFoldGPCR

## Supplementary information

**Supplementary Figure 1.**
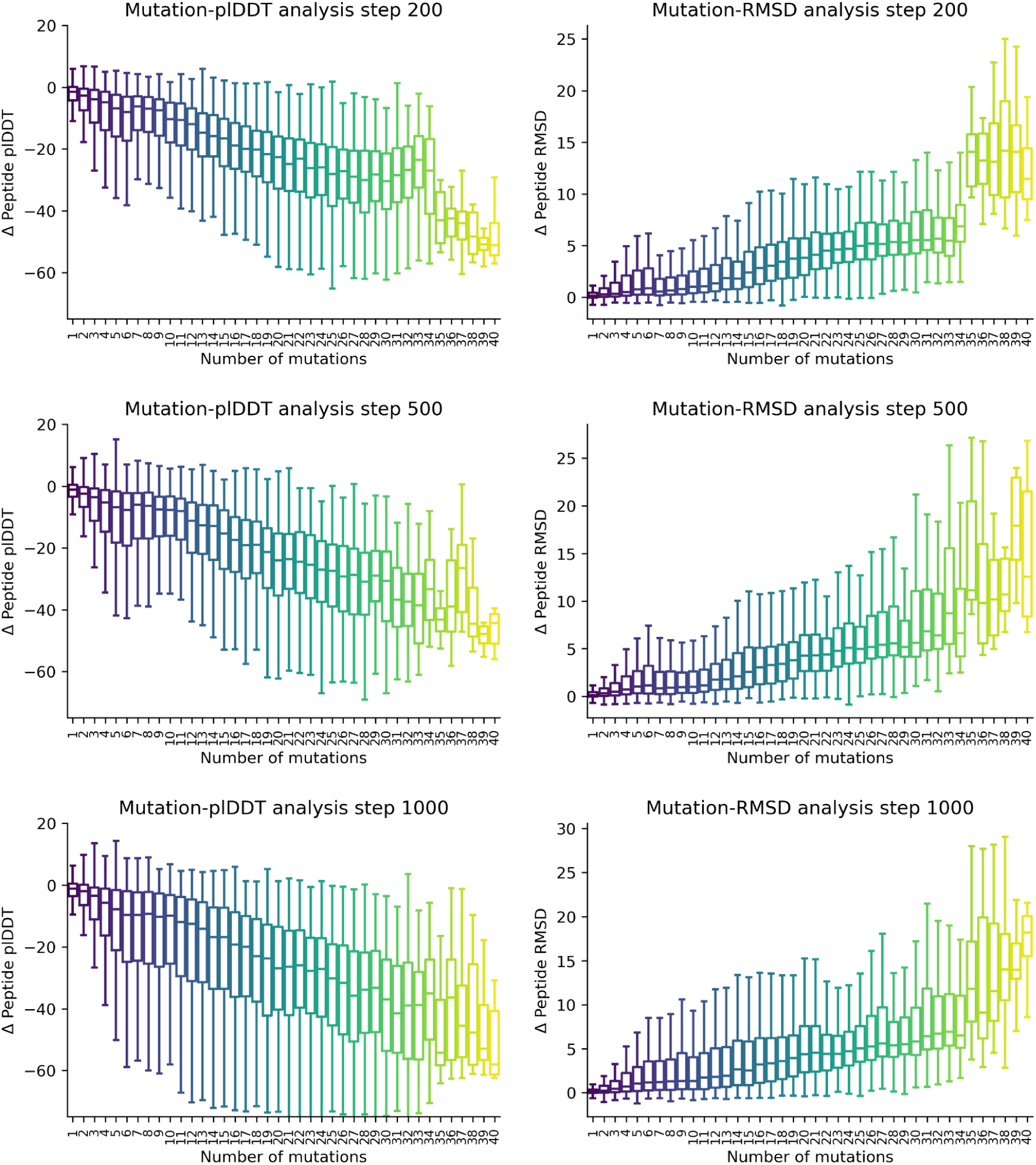
Mutation analysis vs fine-tuning step using the complex model. Peptides with predicted Cβ RMSD < 2 Å were collected at three fine-tuning steps (25’700; 26’000; 26’500). For each set (71, 99, and 126 peptides, respectively), mutations were introduced and analysed as described in Methods.

**Supplementary Figure 2.**
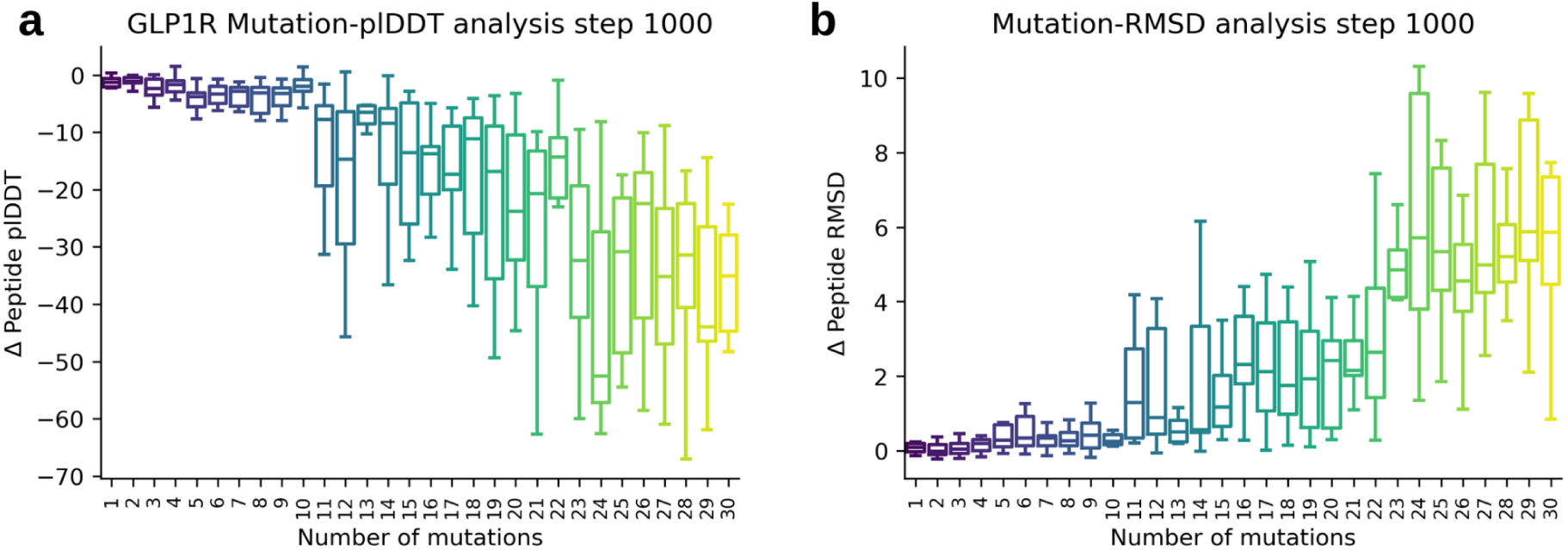
Mutation analysis of GLP1R (PDB ID 6X18, https://www.rcsb.org/structure/6X18) after 1000 fine-tuning steps using the complex model. Ten samples were generated per mutation count. Beyond ∼10 mutations (≈ one-third of the sequence length), model confidence drops sharply and RMSD increases.

**Supplementary Figure 3.**
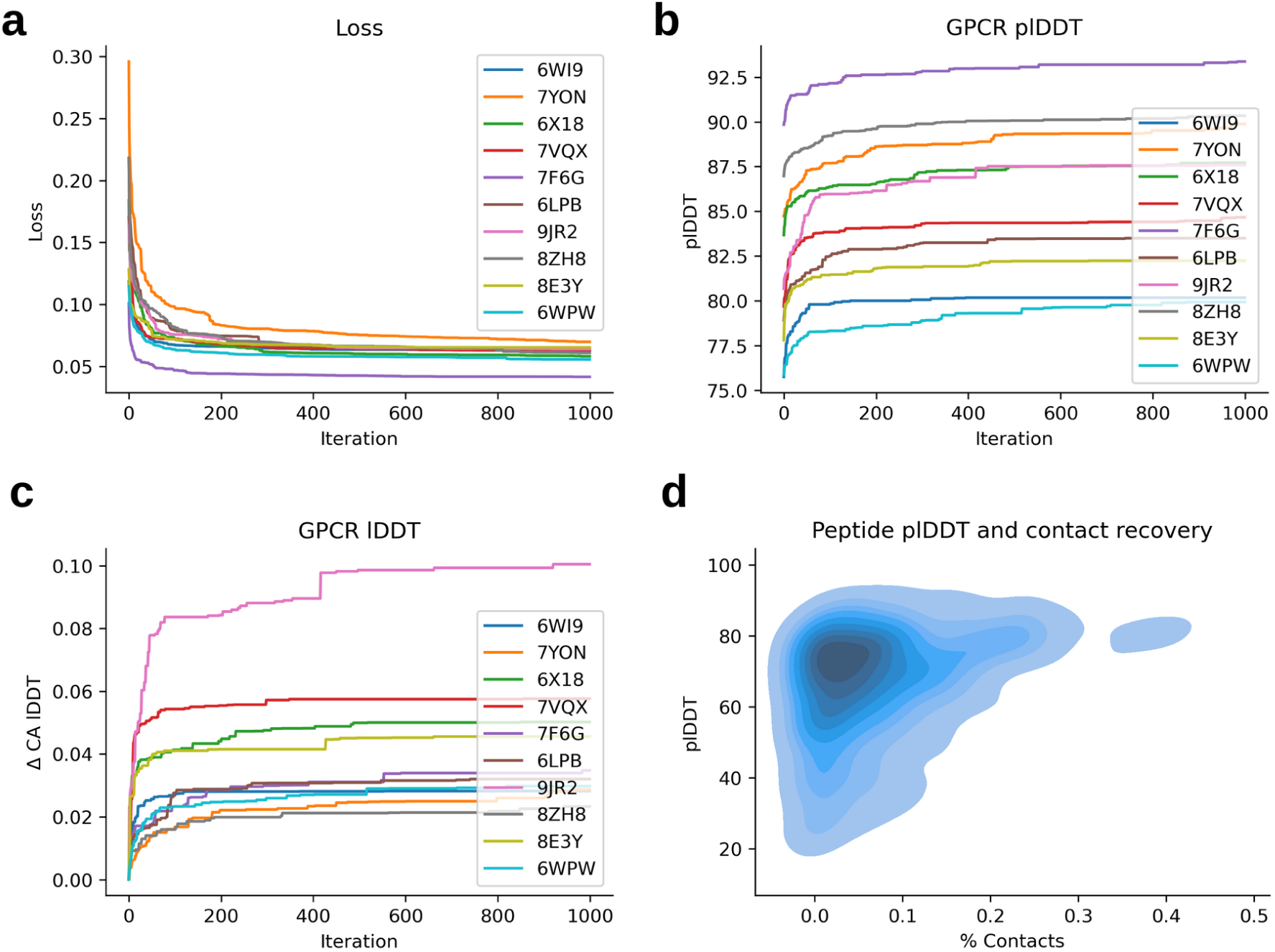
Blind design metrics. Saturation is reached after a few hundred steps in most cases. **a)** Best Loss vs iteration. **b)** Best GPCR plDDT vs iteration. **c)** Best GPCR lDDT. **d)** peptide plDDT vs contact recovery. At high contact recovery, the plDDT is consistently high. However, the plDDT can also be high at lower values, suggesting potential alternative binding modes.

**Supplementary Figure 4.**
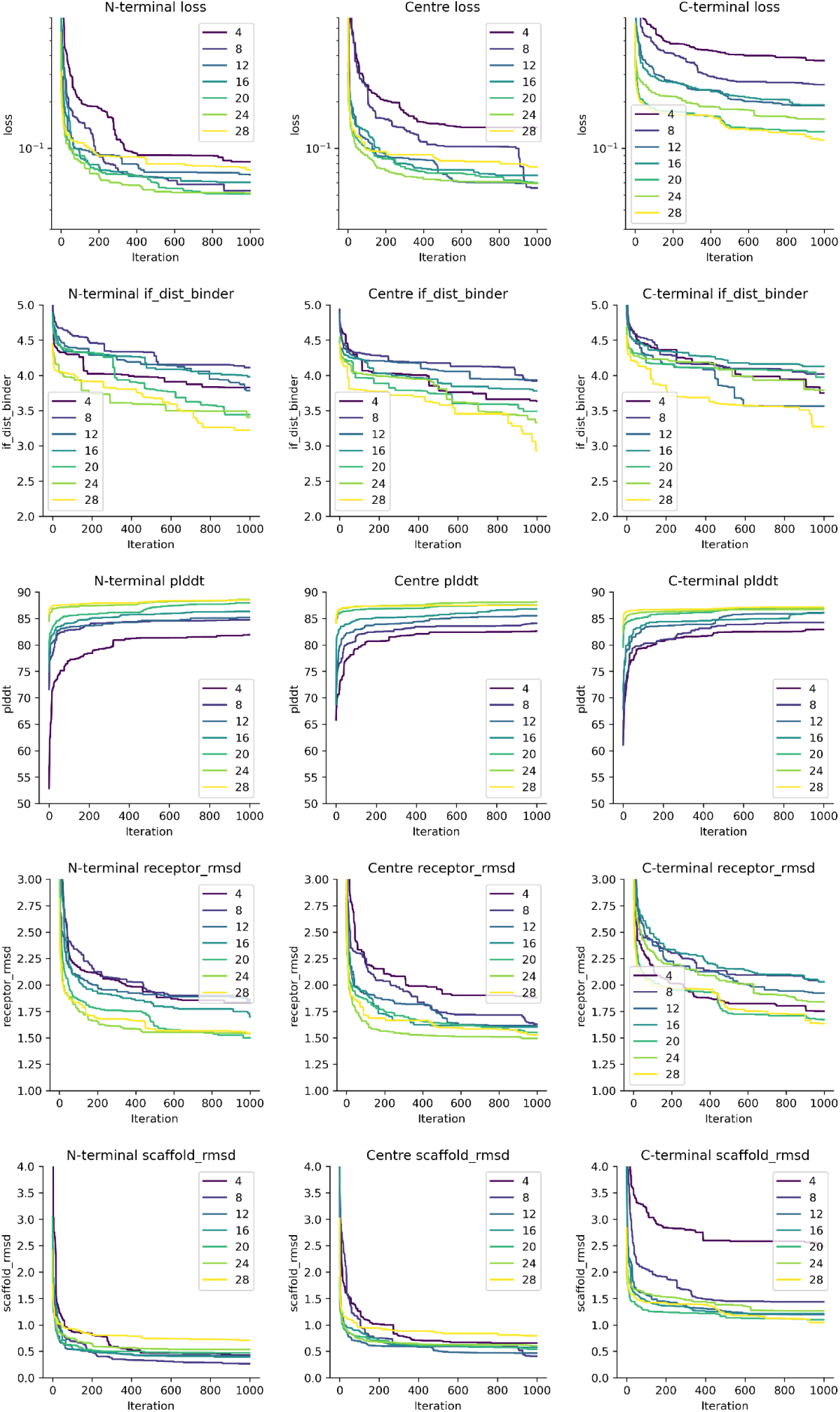
Linear scaffold. Results of linear scaffold designs, divided by scaffold type and coloured by scaffold length. C-terminal scaffolds consistently show higher design losses, likely due to the greater flexibility of the GPCR “lid” in this region, which contrasts with the more rigid N-terminal binding pocket.

**Supplementary Figure 5.**
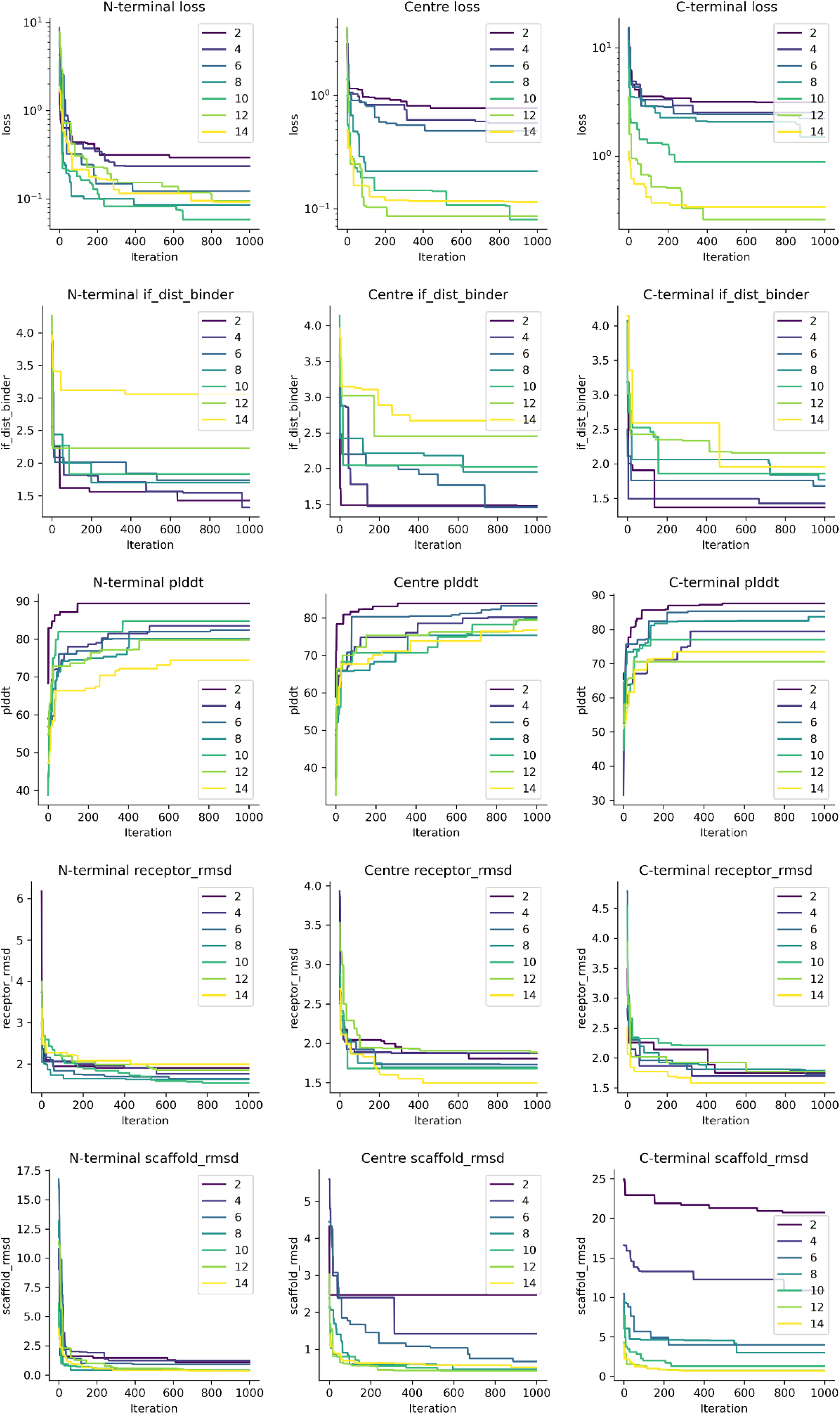
Cyclic scaffold. Longer scaffolding lengths have lower scaffold RMSD, but high plDDT suggesting that even though the scaffold condition can be satisfied, the solution is not predicted to be accurate.

**Supplementary Figure 6.**
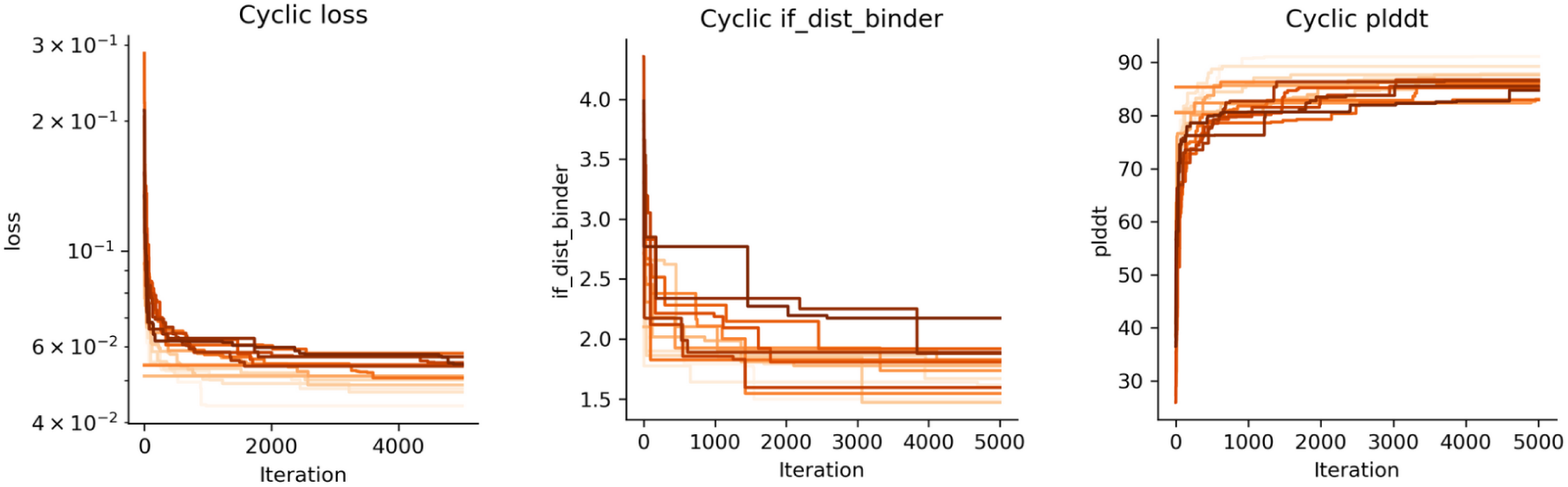
Cyclic blind. Untargeted cyclic peptide design (lengths 15–30 residues). High-confidence cyclic binders were obtained without scaffolding, allowing for the discovery of new binding modes.

**Supplementary Figure 7.**
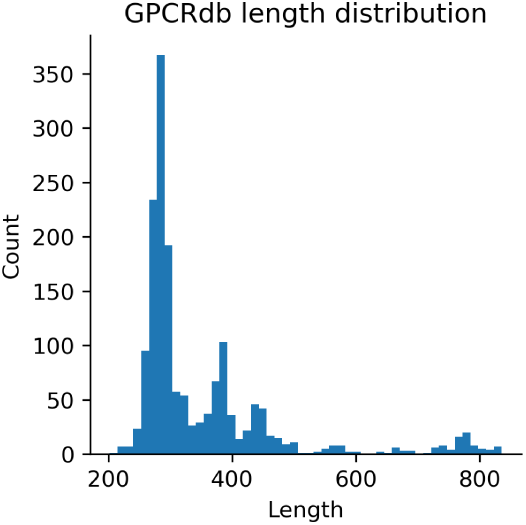
GPCR chain length distribution from single-chain GPCRs in GPCR db (n=1634).

**Supplementary Figure 8.**
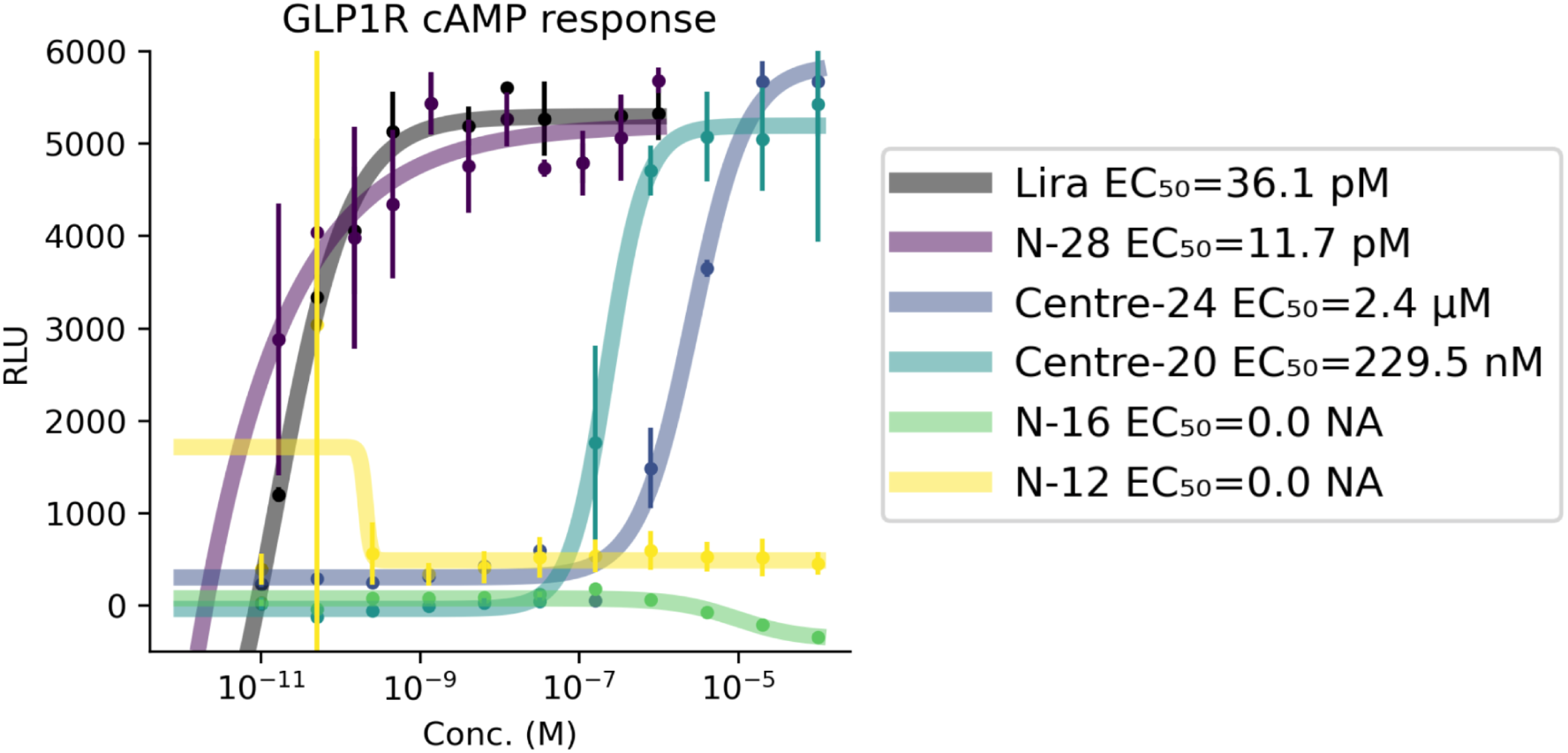
Functional validation for all linear synthesised peptides (RLU=Relative Light Units vs concentration) in GLP1R cAMP cell assay. The EC50s differ slightly from the normalised versions. The lines represent sigmoidal fits, and the points represent averages from three replicates with vertical lines as standards.

**Supplementary Figure 9.**
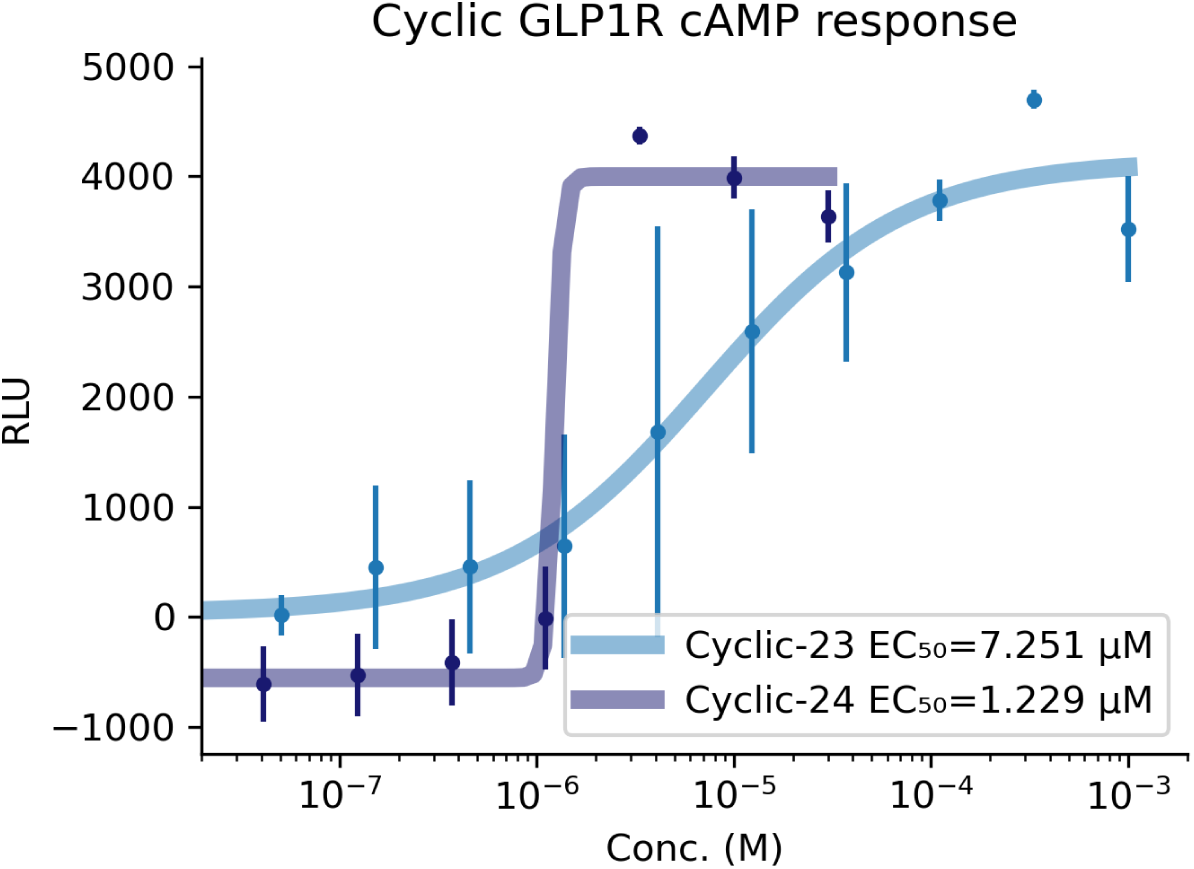
Functional validation for all cyclic synthesised peptides (RLU=Relative Light Units vs concentration) in GLP1R cAMP cell assay. The EC50s differ slightly from the normalised versions. The lines represent sigmoidal fits, and the points represent averages from three replicates with vertical lines as standards.

## One letter sequences

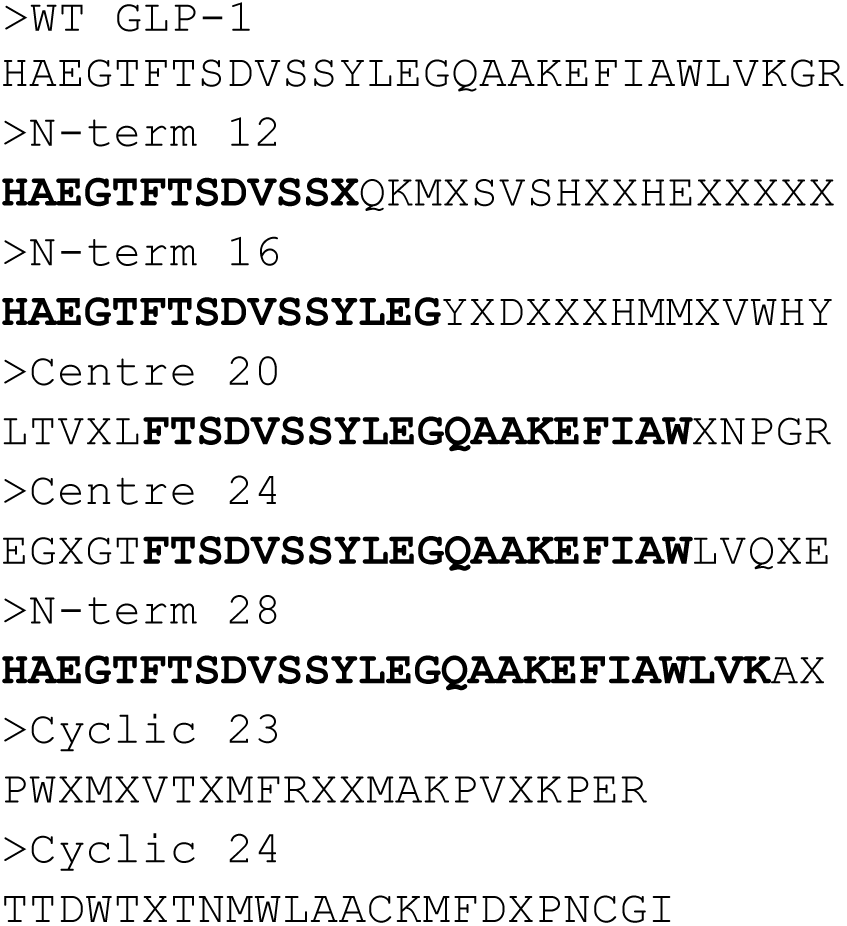

